# Wnt signaling mediates acquisition of blood-brain barrier properties in naïve endothelium derived from human pluripotent stem cells

**DOI:** 10.1101/2021.05.12.443828

**Authors:** Benjamin D. Gastfriend, Hideaki Nishihara, Scott G. Canfield, Koji L. Foreman, Britta Engelhardt, Sean P. Palecek, Eric V. Shusta

**Affiliations:** Department of Chemical and Biological Engineering, University of Wisconsin–Madison, Madison, WI, USA; Theodor Kocher Institute, University of Bern, Bern, Switzerland; Department of Neurological Surgery, University of Wisconsin–Madison, Madison, WI, USA

**Keywords:** Endothelial cells, blood-brain barrier, barriergenesis, Wnt signaling

## Abstract

Endothelial cells (ECs) in the central nervous system (CNS) acquire their specialized blood-brain barrier (BBB) properties in response to extrinsic signals, with Wnt/β-catenin signaling coordinating multiple aspects of this process. Our knowledge of CNS EC development has been advanced largely by animal models, and human pluripotent stem cells (hPSCs) offer the opportunity to examine BBB development in an *in vitro* human system. Here we show that activation of Wnt signaling in hPSC-derived naïve endothelial progenitors, but not in matured ECs, leads to robust acquisition of canonical BBB phenotypes including expression of GLUT-1, increased claudin-5, and decreased PLVAP. RNA-seq revealed a transcriptome profile resembling ECs with CNS-like characteristics, including Wnt-upregulated expression of *LEF1*, *APCDD1*, and *ZIC3*. Together, our work defines effects of Wnt activation in naïve ECs and establishes an improved hPSC-based model for interrogation of CNS barriergenesis.

## INTRODUCTION

In the central nervous system (CNS), vascular endothelial cells (ECs) are highly specialized, with complex tight junctions, expression of a spectrum of nutrient and efflux transporters, low rates of vesicle trafficking, no fenestrae, and low expression of immune cell adhesion molecules (Reese and Karnovsky, 1967; Obermeier et al., 2013). ECs bearing these attributes, often referred to as the blood-brain barrier (BBB), work in concert with the other brain barriers to facilitate the tight regulation of the CNS microenvironment required for proper neuronal function (Daneman and Engelhardt, 2017; Profaci et al., 2020). During development, the Wnt/β-catenin signaling pathway drives both CNS angiogenesis, during which vascular sprouts originating from the perineural vascular plexus invade the developing neural tube, and the coupled process of barriergenesis by which resulting ECs begin to acquire BBB properties (Liebner et al., 2008; Stenman et al., 2008; Daneman et al., 2009; Engelhardt and Liebner, 2014; Umans et al., 2017). Specifically, neural progenitor-derived Wnt7a and Wnt7b ligands signal through Frizzled receptors and the obligate co-receptors RECK and GPR124 (ADGRA2) on endothelial cells (Kuhnert et al., 2010; Cullen et al., 2011; Vanhollebeke et al., 2015; Cho et al., 2017; Eubelen et al., 2018; Vallon et al., 2018). Other ligands function analogously in the retina (Norrin) (Ye et al., 2009; Wang et al., 2012) and potentially in the dorsal neural tube (Daneman et al., 2009). Furthermore, Wnt/β-catenin signaling is required for maintenance of CNS EC barrier properties in adulthood (Tran et al., 2016), with astrocytes as a major source of Wnt7 ligands (He et al., 2018; Vanlandewijck et al., 2018; Guérit et al., 2021).

Molecular hallmarks of Wnt-mediated CNS EC barriergenesis are (i) acquisition of glucose transporter GLUT-1 expression, (ii) loss of plasmalemma vesicle-associated protein (PLVAP), and (iii) upregulation of claudin-5 (Daneman et al., 2009; Kuhnert et al., 2010; Cho et al., 2017; Umans et al., 2017; Wang et al., 2019). Notably, the Wnt-mediated switch between the “leaky” EC phenotype (GLUT-1^-^ PLVAP^+^ claudin-5^low^) and the barrier EC phenotype (GLUT-1^+^ PLVAP^-^ claudin-5^high^) correlates with reduced permeability to molecular tracers (Wang et al., 2012; Cho et al., 2017) and is conserved in multiple contexts. For instance, medulloblastomas that produce Wnt-inhibitory factors have leaky vessels (Phoenix et al., 2016). Moreover, vasculature perfusing circumventricular organs is leaky due to low levels of Wnt signaling (Benz et al., 2019; Wang et al., 2019). Notably, ectopic activation of Wnt in ECs of circumventricular organs induces GLUT-1 and suppresses PLVAP (Benz et al., 2019; Wang et al., 2019). However, similar ectopic activation of Wnt in liver and lung ECs produces only very minor barriergenic effects (Munji et al., 2019), and Wnt activation in cultured primary mouse brain ECs does not prevent culture-induced loss of barrier-associated gene expression (Sabbagh and Nathans, 2020). The reasons for the apparent context-dependent impacts of Wnt activation in ECs remain unclear and motivate systematic examination of this process in a simplified model system. Further, given species differences in brain EC transporter expression (Uchida et al., 2011), drug permeability (Syvänen et al., 2009), and gene expression (Song et al., 2020), this process warrants investigation in human cells to complement mouse *in vivo* studies.

Prior studies have evaluated the impact of Wnt activation in immortalized human brain ECs and observed only modest effects on barrier phenotype (Paolinelli et al., 2013; Laksitorini et al., 2019). Combined with the aforementioned deficits observed in primary adult mouse brain endothelial cells that are not rescued by ectopic Wnt activation (Sabbagh and Nathans, 2020), one possibility is that mature, adult endothelium is largely refractory to Wnt activation, and that Wnt responsiveness is a property of immature endothelial cells analogous to those in the perineural vascular plexus. Human pluripotent stem cells (hPSCs) offer a potential human model system for investigation of molecular mechanisms of BBB phenotype acquisition. However, currently available hPSC-based models of CNS endothelial-like cells are not well suited for modeling the BBB developmental progression as they do not follow a developmentally-relevant differentiation trajectory, lack definitive endothelial identity, or have been incompletely characterized with respect to the role of developmental signaling pathways (Lippmann et al., 2020; Workman and Svendsen, 2020). As a potential alternative, hPSCs can also be used to generate immature, naïve endothelial progenitors (Lian et al., 2014) that could be used to better explore the induction of BBB phenotypes. For example, we recently reported that extended culture of such hPSC-derived endothelial progenitors in a minimal medium yielded ECs with improved BBB tight junction protein expression and localization which led to improved paracellular barrier properties (Nishihara et al., 2020). However, as shown below, these cells exhibit high expression of PLVAP and little expression of GLUT-1, indicating the need for additional cues to drive CNS EC specification.

In this work, we tested the hypothesis that activation of Wnt/β-catenin signaling in hPSC-derived, naïve endothelial progenitors would drive development of a CNS EC-like phenotype. We found that many aspects of the CNS EC phenotype, including the canonical GLUT-1, claudin-5, and PLVAP expression effects, were regulated by CHIR 99021, a small molecule agonist of Wnt/β-catenin signaling. Wnt ligands and conditioned media from neural progenitors produced a more limited response, as did CHIR treatment in matured ECs. Whole-transcriptome analysis revealed definitive endothelial identity of the resulting cells and CHIR-upregulated expression of known CNS EC transcripts, including *LEF1*, *APCDD1*, *AXIN2*, *SLC2A1*, *CLDN5*, *LSR*, *ABCG2*, *SOX7*, and *ZIC3*. We also observed an unexpected CHIR-mediated upregulation of caveolin-1, which did not, however, correlate with increased uptake of a dextran tracer. Thus, we provide evidence that Wnt activation in hPSC-derived naïve endothelial progenitors is sufficient to induce many aspects of the CNS barrier EC phenotype, and we establish a model system for further systematic investigation of putative barriergenic cues.

## RESULTS

### Wnt activation in hPSC-derived endothelial progenitors

We adapted an existing protocol to produce endothelial progenitor cells (EPCs) from hPSCs (Lian et al., 2014; Bao et al., 2016) (Figure 1A). To achieve mesoderm specification, this method employs an initial activation of Wnt/β-catenin signaling with CHIR 99021 (CHIR), a small molecule inhibitor of glycogen synthase kinase-3 (GSK-3), which results in inhibition of GSK-3β-mediated β-catenin degradation. After 5 days of expansion, the resulting cultures contained a mixed population of CD34^+^CD31^+^ EPCs and CD34^-^CD31^-^ non-EPCs (Figure 1B-C). We used magnetic-activated cell sorting (MACS) to isolate CD31^+^ cells from this mixed culture and plated these cells on collagen IV-coated plates in a minimal endothelial cell medium termed hECSR (Nishihara et al., 2020). We first asked whether Wnt3a, a ligand widely used to activate canonical Wnt/β-catenin signaling (Kim et al., 2005, 2008; Liebner et al., 2008; Cecchelli et al., 2014; Praça et al., 2019), could induce GLUT-1 expression in the resulting ECs. After 6 days of treatment, we observed a significant increase in the fraction of GLUT-1^+^ ECs in Wnt3a-treated cultures compared to controls (Figure 1D-E). Consistent with previous observations (Nishihara et al., 2020), we also detected a population of calponin^+^ smooth muscle protein 22-⍺^+^ putative smooth muscle-like cells (SMLCs) outside the endothelial colonies (Figure 1–figure supplement 1) and these SMLCs expressed GLUT-1 in both control and Wnt3a-treated conditions (Figure 1D).

**Figure 1.**
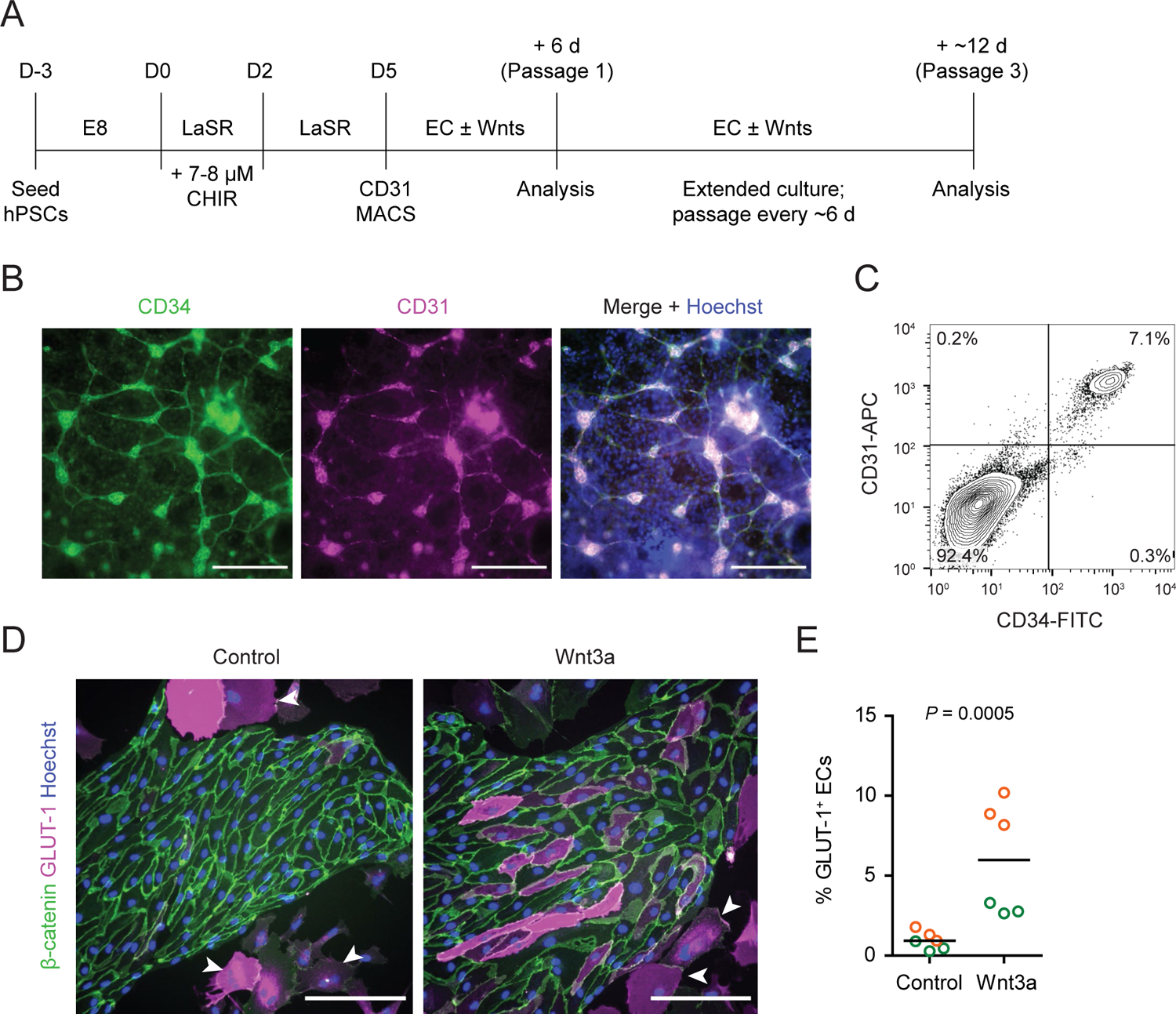
hPSC-derived endothelial progenitors as a model for studying Wnt-mediated barriergenesis. (A) Overview of the endothelial differentiation and Wnt treatment protocol. **(B)** Immunocytochemistry analysis of CD34 and CD31 expression in D5 EPCs prior to MACS. Hoechst nuclear counterstain is overlaid in the merged image. Scale bars: 200 μm. **(C)** Flow cytometry analysis of CD34 and CD31 expression in D5 EPCs prior to MACS. **(D)** Immunocytochemistry analysis of β-catenin and GLUT-1 expression in Passage 1 ECs treated with Wnt3a or control. Hoechst nuclear counterstain is overlaid. Arrowheads indicate smooth muscle-like cells (SMLCs). Scale bars: 200 μm. **(E)** Quantification of the percentage of GLUT-1^+^ ECs in control- and Wnt3a-treated conditions. Points represent replicate wells from 2 independent differentiations of the IMR90-4 line, each differentiation indicated with a different color. Bars indicate mean values. P-value: Two-way ANOVA.

**Figure 1–figure supplement 1.**
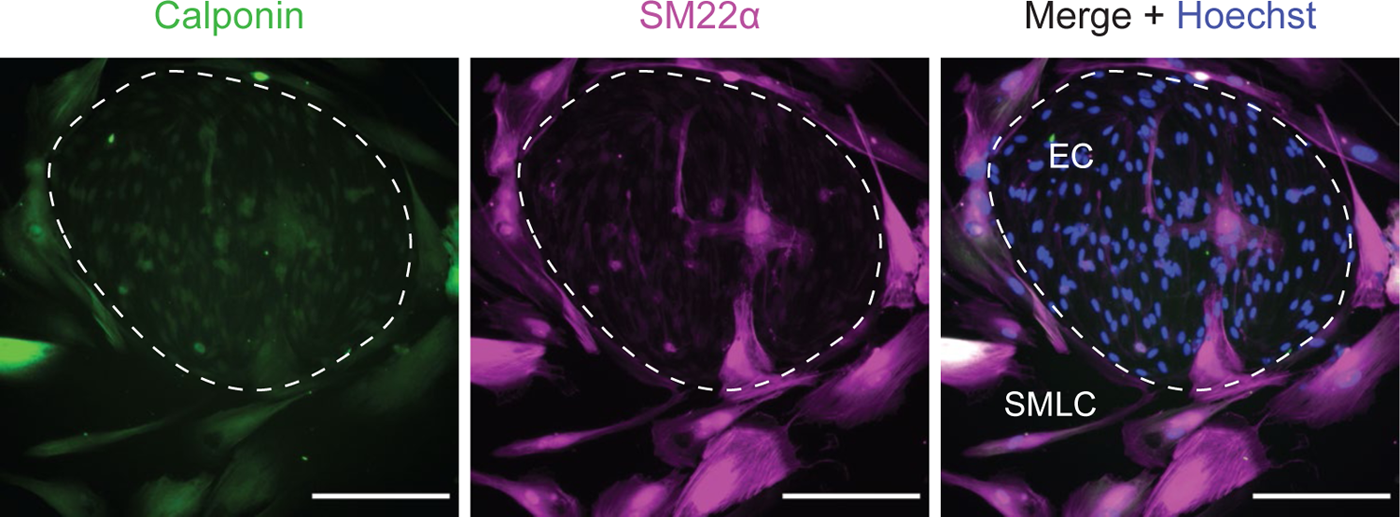
Smooth-muscle like cells (SMLCs). Immunocytochemistry analysis of calponin and smooth muscle protein 22-⍺ (SM22⍺) in Passage 1 cultures containing ECs and SMLCs. Hoechst nuclear counterstain is overlaid in the merged image. Dashed area indicates an EC colony. Scale bars: 200 μm.

Based on these promising results with Wnt3a, we next tested several additional strategies for Wnt activation and, in addition to GLUT-1, evaluated expression of two other key proteins: claudin-5, which is known to be upregulated in CNS ECs in response to Wnt (Benz et al., 2019), and caveolin-1, given the low rate of caveolin-mediated transcytosis in CNS compared to non-CNS ECs (Reese and Karnovsky, 1967; Andreone et al., 2017). First, we tested Wnt7a and Wnt7b, the ligands primarily responsible for Wnt activation in CNS ECs *in vivo* (Daneman et al., 2009; Cho et al., 2017). We also tested Wnt ligands in combination with R-spondin 1 (Rspo1), a potentiator of Wnt signaling that inhibits the RNF43/ZNRF3-mediated negative feedback mechanism by which Frizzled receptors are endocytosed (Kim et al., 2005, 2008; Koo et al., 2012; Clevers et al., 2014). Finally, we tested a low concentration (4 µM) of the GSK-3 inhibitor CHIR because of its ability to activate Wnt signaling in a receptor/co-receptor-independent manner. We found that Wnt7a and the combination of Wnt7a and Wnt7b, but not Wnt7b alone, slightly increased the fraction of GLUT-1^+^ ECs, while Rspo1 did not affect EC purity or expression of GLUT-1, claudin-5 or caveolin-1 (Figure 2A-C). Interestingly, Wnt7a, but not Wnt3a, also increased the proportion of ECs compared to SMLCs (Figure 2A,C). By contrast, 4 µM CHIR robustly induced GLUT-1 expression in approximately 90% of ECs while increasing EC purity to a level similar to that achieved with Wnt7a. Furthermore, CHIR led to an approximately 1.5-fold increase in average claudin-5 abundance and a nearly 30-fold increase in GLUT-1 abundance, but also a 4-fold increase in caveolin-1 (Figure 2A,C). We therefore titrated CHIR to determine an optimal concentration for EC expansion, purity, GLUT-1 induction, and claudin-5 upregulation while limiting the undesirable non-CNS-like increase in caveolin-1 abundance. Although 2 µM CHIR did not lead to increased caveolin-1 expression compared to vehicle control (DMSO), it also did not elevate claudin-5 or GLUT-1 expression compared to control and was less effective in increasing EC number and EC purity than 4 µM CHIR (Figure 2–figure supplement 1). On the other hand, 6 µM CHIR further increased GLUT-1 abundance but also further increased caveolin-1 abundance and did not improve EC number, EC purity, or claudin-5 expression (Figure 2–figure supplement 1). Therefore, we conducted further experiments using 4 µM CHIR. We confirmed that the CHIR-mediated increases in EC purity, EC number, and caveolin-1 and GLUT-1 expression were conserved in an additional hPSC line, although claudin-5 upregulation was not apparent (Figure 2–figure supplement 2). We also used two hPSC lines with doxycycline-inducible expression of short hairpin RNAs targeting *CTNNB1* (β-catenin) to confirm that CHIR-mediated upregulation of GLUT-1 in ECs was β-catenin-dependent. Indeed, doxycycline treatment in combination with CHIR significantly reduced GLUT-1 abundance in ECs derived from these hPSC lines (Figure 2–figure supplement 3). Together, these results suggest that Wnt pathway activation, either with ligands or CHIR, is capable of inducing CNS-like phenotypes in hPSC-derived endothelial progenitors.

**Figure 2.**
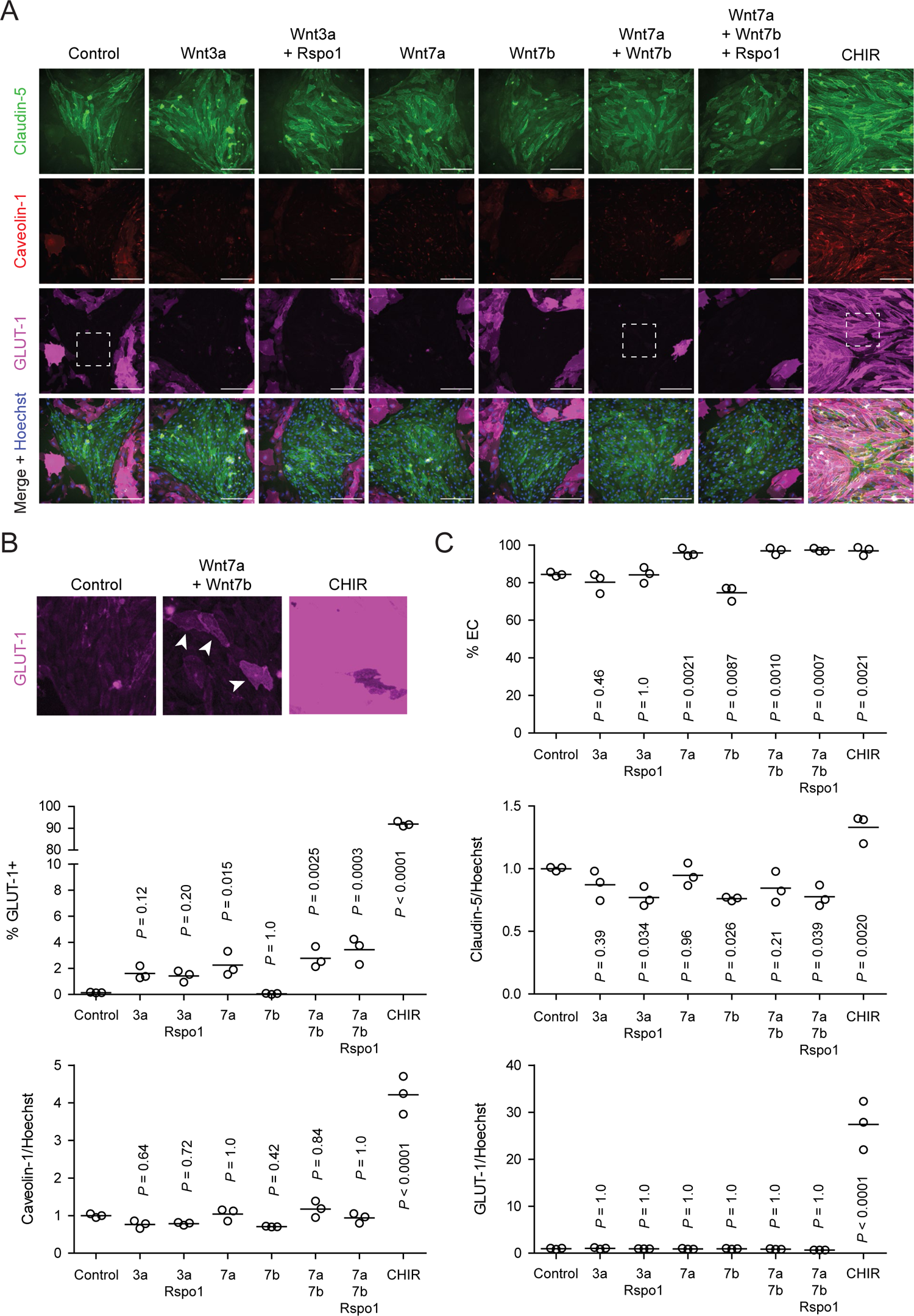
Effect of Wnt ligands and pathway modulators on endothelial properties. **(A)** Immunocytochemistry analysis of claudin-5, caveolin-1, and GLUT-1 expression in Passage 1 ECs treated with Wnt3a, Wnt3a + R-spondin 1 (Rspo1), Wnt7a, Wnt7b, Wnt7a + Wnt7b, Wnt7a + Wnt7b + Rspo1, CHIR, or control. Hoechst nuclear counterstain is overlaid in the merged images. Dashed boxes indicate fields displayed in (B). Scale bars: 200 μm. **(B)** Immunocytochemistry analysis of GLUT-1 expression in the fields indicated with dashed boxes in (A) from the control and Wnt7a + Wnt7b conditions. To visualize weak GLUT-1 immunoreactivity in Wnt7a + Wnt7b-treated ECs, a linear brightness/contrast adjustment was applied identically to the three fields but differs from that of the images shown in (A). Arrowheads indicate GLUT-1^+^ ECs. **(C)** Quantification of images from the conditions described in (A) for percentage of ECs (claudin-5^+^ cells relative to total nuclei), GLUT-1^+^ ECs (relative to total claudin-5^+^ ECs), and mean fluorescence intensity of claudin-5, caveolin-1, and GLUT-1 normalized to Hoechst mean fluorescence intensity within the area of claudin-5^+^ ECs only. Points represent replicate wells from one differentiation of the IMR90-4 line and bars indicate mean values. For the fluorescence intensity plots, values were normalized such that the mean of the control condition equals 1. P-values: ANOVA followed by Dunnett’s test versus control.

**Figure 2–figure supplement 1.**
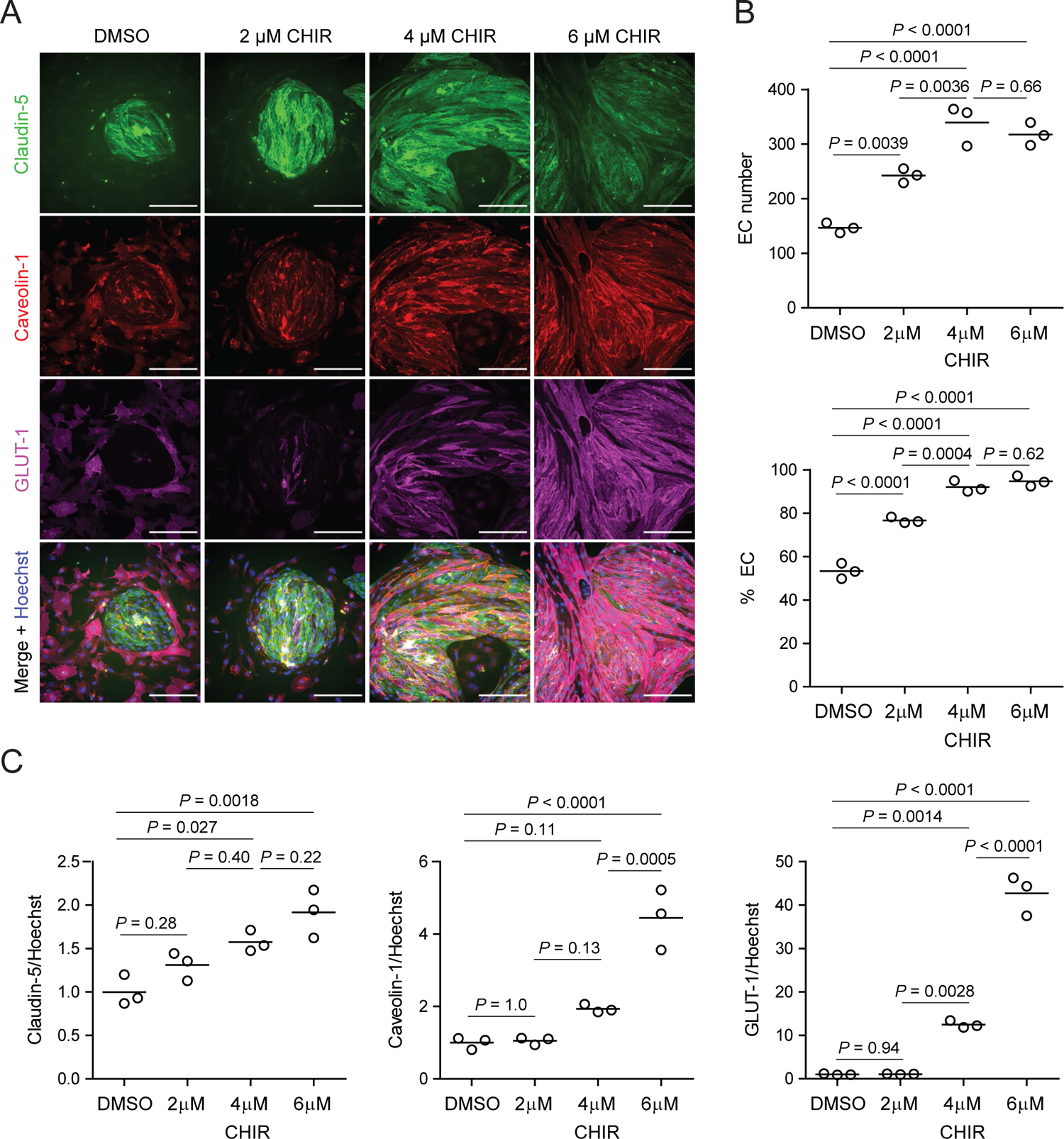
Dose-dependent effects of CHIR on endothelial properties. **(A)** Immunocytochemistry analysis of claudin-5, caveolin-1, and GLUT-1 expression in Passage 1 ECs treated with 2 μM, 4 μM, or 6 μM CHIR, or DMSO vehicle control. Hoechst nuclear counterstain is overlaid in the merged images. Scale bars: 200 μm. **(B)** Quantification of images from the conditions described in (A) for number of ECs per 20× field and percentage of ECs (claudin-5^+^ cells relative to total nuclei). Points represent replicate wells from one differentiation of the IMR90-4 line and bars indicate mean values. P-values: ANOVA followed by Tukey’s HSD test. **(C)** Quantification of claudin-5, caveolin-1, and GLUT-1 mean fluorescence intensity normalized to Hoechst mean fluorescence intensity within the area of claudin-5^+^ ECs only. Points represent replicate wells from one differentiation of the IMR90-4 line. Bars indicate mean values, with values normalized such that the mean of the DMSO condition equals 1. P-values: ANOVA followed by Tukey’s HSD test.

**Figure 2–figure supplement 2.**
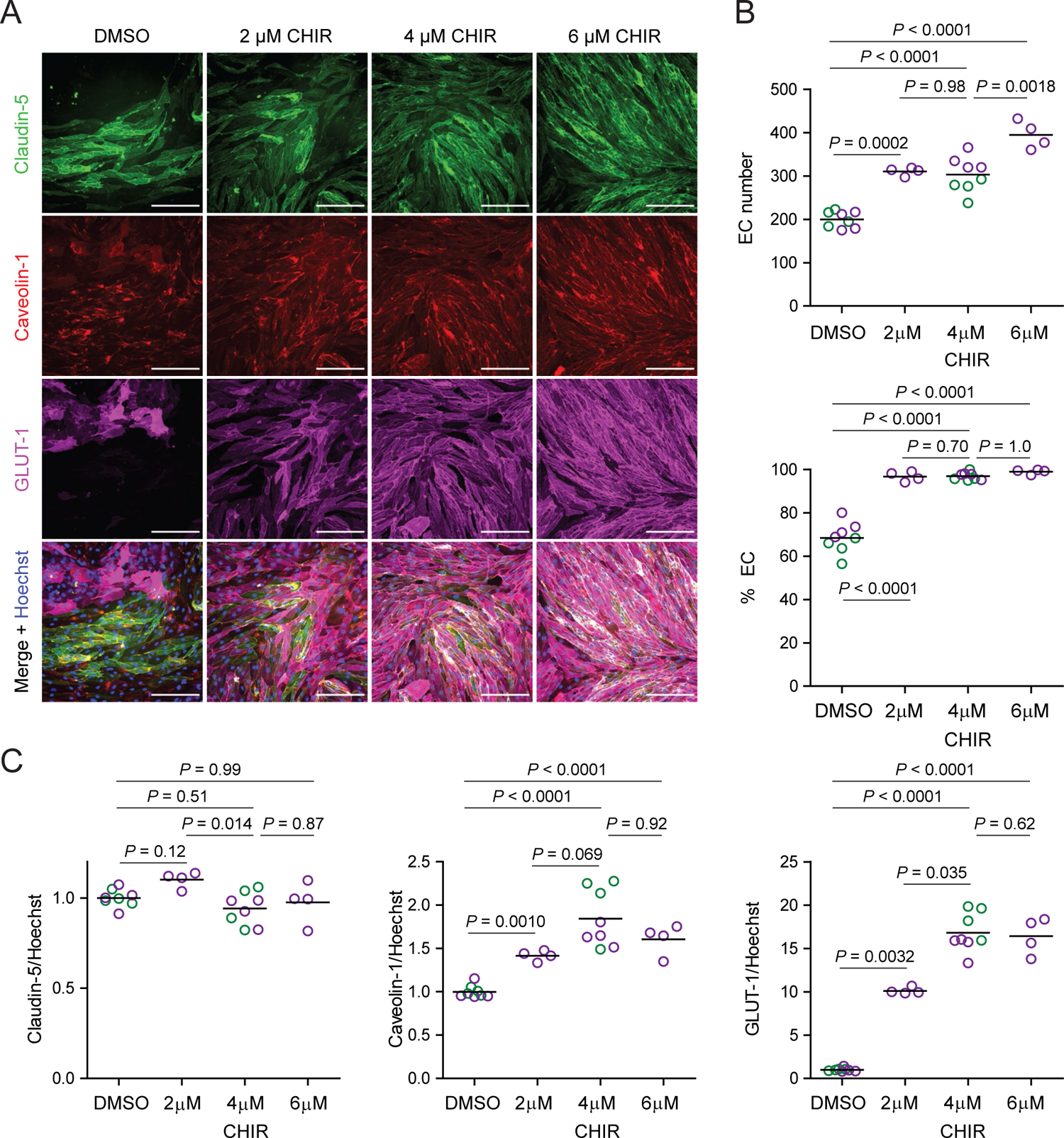
CHIR-mediated effects in an additional hPSC line. **(A)** Immunocytochemistry analysis of claudin-5, caveolin-1, and GLUT-1 expression in Passage 1 ECs differentiated from the WTC11 iPSC line treated with 2 μM, 4 μM, or 6 μM CHIR, or DMSO vehicle control. Hoechst nuclear counterstain is overlaid in the merged images. Scale bars: 200 μm. **(B)** Quantification of images from the conditions described in (A) for number of ECs per 20× field and percentage of ECs (claudin-5^+^ cells relative to total nuclei). Points represent replicate wells from 1–2 differentiations of the WTC11 line and bars indicate mean values, each differentiation indicated with a different color. P-values: Two-way ANOVA followed by Tukey’s HSD test. **(C)** Quantification of claudin-5, caveolin-1, and GLUT-1 mean fluorescence intensity normalized to Hoechst mean fluorescence intensity within the area of claudin-5^+^ ECs only. Points represent replicate wells from 1–2 differentiations of the WTC11 line, each differentiation indicated with a different color. Bars indicate mean values, with values normalized within each differentiation such that the mean of the DMSO condition equals 1. P-values: Two-way ANOVA followed by Tukey’s HSD test on unnormalized data.

**Figure 2–figure supplement 3.**
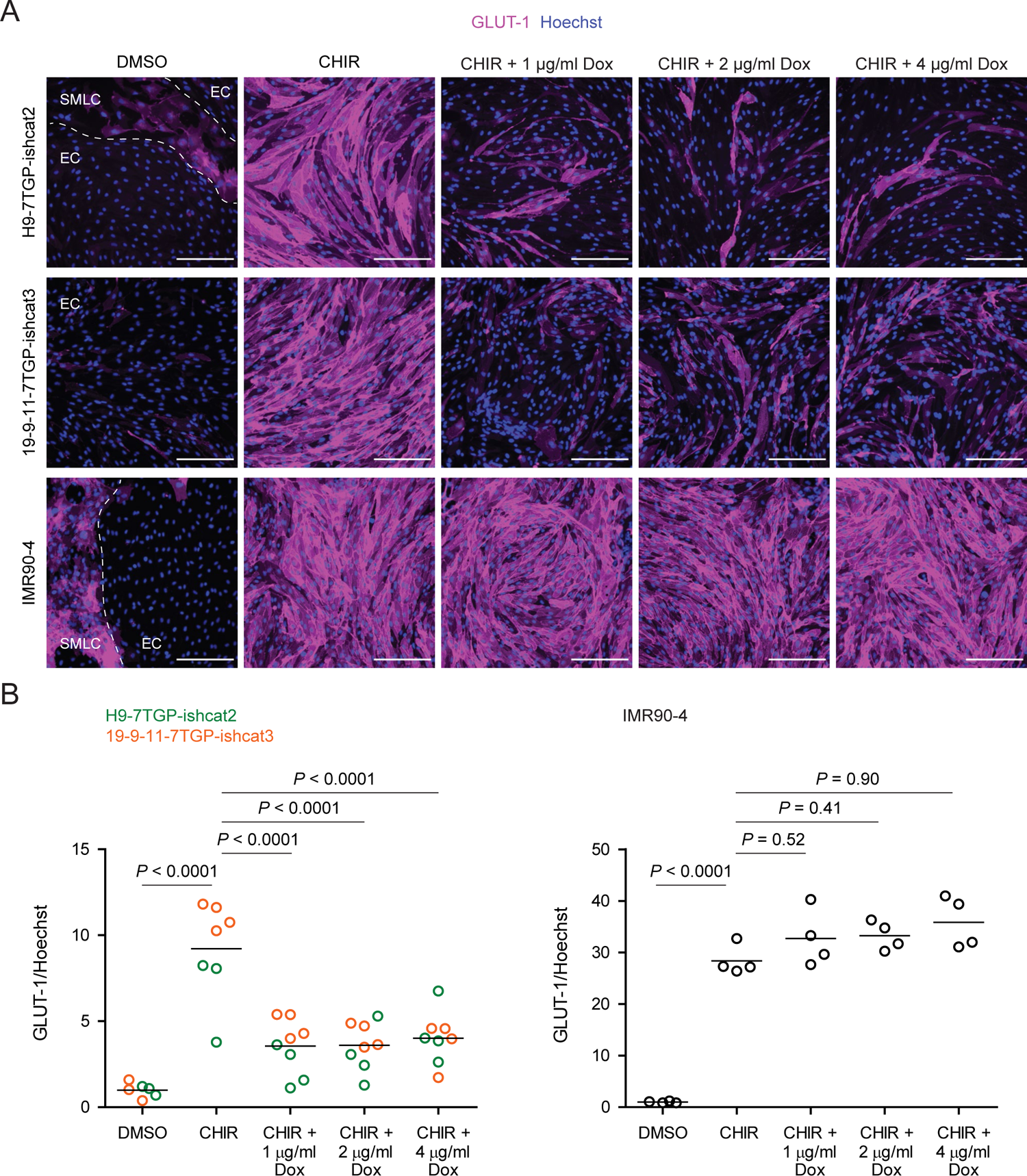
β-catenin-dependence of CHIR-mediated GLUT-1 induction. **(A)** Immunocytochemistry analysis of GLUT-1 expression in Passage 1 ECs treated with DMSO, CHIR, or CHIR + doxycycline (Dox) at 1, 2, or 4 μg/ml. Images from the H9-7TGP-ishcat2, 19-9-11-7TGP-ishcat3, and IMR90-4 lines are shown. Hoechst nuclear counterstain is overlaid. Dashed lines indicate borders between EC colonies and SMLCs in the DMSO condition. Scale bars: 200 μm. **(B)** Quantification of images from the conditions described in (A) for GLUT-1 mean fluorescence intensity normalized to Hoechst mean fluorescence intensity within the area of ECs only. At left, points represent replicate wells from one differentiation of the H9-7TGP-ishcat line (green) and one differentiation of the 19-9-11-7TGP-ishcat3 line (orange). Bars indicate mean values, with values normalized within each differentiation such that the mean of the DMSO condition equals 1. P-values: Two-way ANOVA followed by Tukey’s HSD test on unnormalized data. At right, points represent replicate wells from one differentiation of the IMR90-4 line. Bars indicate mean values, with values normalized such that the mean of the DMSO condition equals 1. P-values: ANOVA followed by Tukey’s HSD test.

In the CNS, neural progenitors and astrocytes are the primary sources of Wnt ligands resulting in induction and maintenance of EC barrier properties. Because the relatively weak response to Wnt ligands observed in our system is potentially attributable to poor potency associated with the recombinant proteins, we reasoned that relevant cellular sources of Wnt ligands might be more effective in activating Wnt in EPCs. To this end, we differentiated hPSCs to neural rosettes, which are radially organized Pax6^+^ neural progenitors, and astrocytes according to established protocols (Ebert et al., 2013; Lippmann et al., 2014; Sareen et al., 2014; Canfield et al., 2017). Importantly, RNA-seq data from the literature suggest that both hPSC-derived neural rosettes and astrocytes express *WNT7A* (Vatine et al., 2016; Shang et al., 2018). We collected neural rosette-conditioned medium (NR-CM) and astrocyte-conditioned medium (Astro-CM) and treated EPCs with these media for 6 days. Similar to our observations with Wnt7a, both NR-CM and Astro-CM significantly increased the proportion of ECs compared to SMLCs (Figure 3A,C). NR-CM, but not Astro-CM, also induced weak GLUT-1 expression in ECs, reminiscent of the Wnt7a-induced phenotype, although this induction was much weaker than in the CHIR-treated cells (Figure 3B,D). NR-CM and Astro-CM had variable effects with respect to caveolin-1 and claudin-5 expression (Figure 3D). In summary, NR-CM performed similarly to Wnt7a in weakly inducing GLUT-1 expression and increasing EC purity. The comparatively stronger response to CHIR may suggest either that the potency or concentration of ligands is insufficient, or that the EPCs lack the full machinery of receptors and co-receptors necessary to transduce the Wnt ligand signal (analyzed further below).

**Figure 3.**
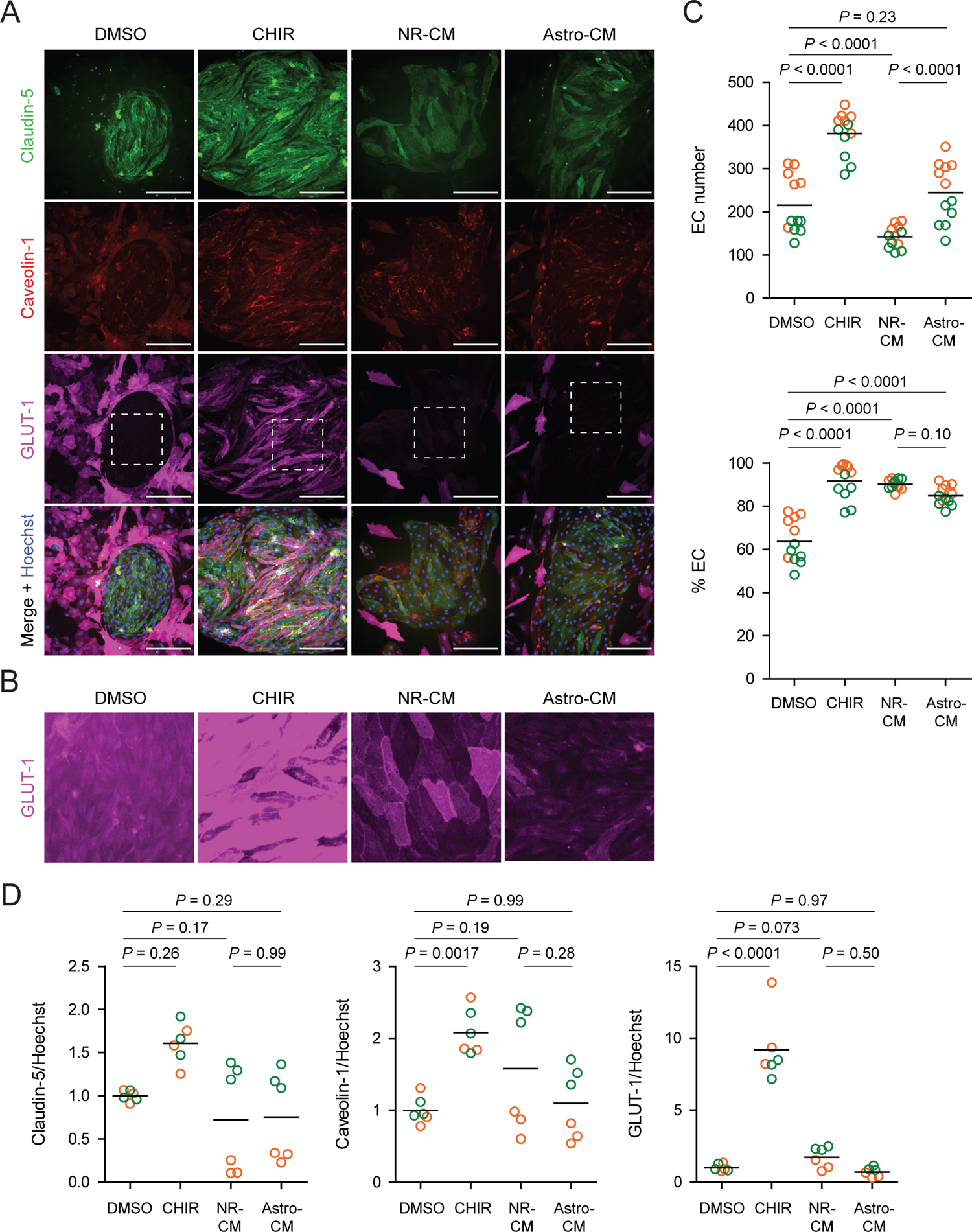
Effect of neural rosette- and astrocyte-conditioned media on endothelial properties. (A) Immunocytochemistry analysis of claudin-5, caveolin-1, and GLUT-1 expression in Passage 1 ECs treated with DMSO, CHIR, neural rosette-conditioned medium (NR-CM), or astrocyte-conditioned medium (Astro-CM). Hoechst nuclear counterstain is overlaid in the merged images. Dashed boxes indicate fields displayed in (B). Scale bars: 200 μm. **(B)** Immunocytochemistry analysis of GLUT-1 expression in the fields indicated with dashed boxes in (A). A linear brightness/contrast adjustment was applied identically to the four fields but differs from that of the images shown in (A). **(C)** Quantification of images from the conditions described in (A) for number of ECs per 20× field and percentage of ECs (claudin-5^+^ cells relative to total nuclei). Points represent replicate wells from two independent differentiations of the IMR90-4 line, each differentiation indicated with a different color. Bars indicate mean values. P-values: Two-way ANOVA followed by Tukey’s HSD test. **(D)** Quantification of claudin-5, caveolin-1, and GLUT-1 mean fluorescence intensity normalized to Hoechst mean fluorescence intensity within the area of claudin-5^+^ ECs only. Points represent replicate wells from two independent differentiations of the IMR90-4 line, each differentiation indicated with a different color. Bars indicate mean values, with values normalized within each differentiation such that the mean of the DMSO condition equals 1. P-values: Two-way ANOVA followed by Tukey’s HSD test on unnormalized data.

### Effects of CHIR-mediated Wnt activation in endothelial progenitors

Since CHIR elicited the most robust Wnt-mediated response, we next asked whether other aspects of the CNS EC barrier phenotype were CHIR-regulated. PLVAP, a protein that forms bridges across both caveolae and fenestrae (Herrnberger et al., 2012), is one such canonically Wnt-downregulated protein. We therefore first evaluated PLVAP expression in Passage 1 control (DMSO) or CHIR-treated ECs using confocal microscopy (Figure 4A). We observed numerous PLVAP^+^ punctate vesicle-like structures in both conditions, with CHIR treatment reducing PLVAP abundance by approximately 20% (Figure 4A-B). This effect was not apparent in Western blots of Passage 1 ECs, likely due to the relatively modest effect (Figure 5A-B). However, after two more passages (Figure 1A), Passage 3 ECs demonstrated a robust downregulation of PLVAP in CHIR-treated cells compared to controls (Figure 5C-D). We also used Western blotting to confirm CHIR-mediated upregulation of GLUT-1 and claudin-5 both at Passage 1 and Passage 3 (Figure 5A-D). We next evaluated expression of the tricellular tight junction protein LSR (angulin-1) because of its enrichment in CNS versus non-CNS ECs, and the temporal similarity between LSR induction and the early stage of Wnt-mediated CNS barriergenesis (Sohet et al., 2015). We found that CHIR treatment led to a strong increase in LSR expression in both Passage 1 and Passage 3 ECs (Figure 5A-D), suggesting that Wnt signaling upregulates multiple necessary components of the CNS EC bicellular and tricellular junctions.

**Figure 4.**
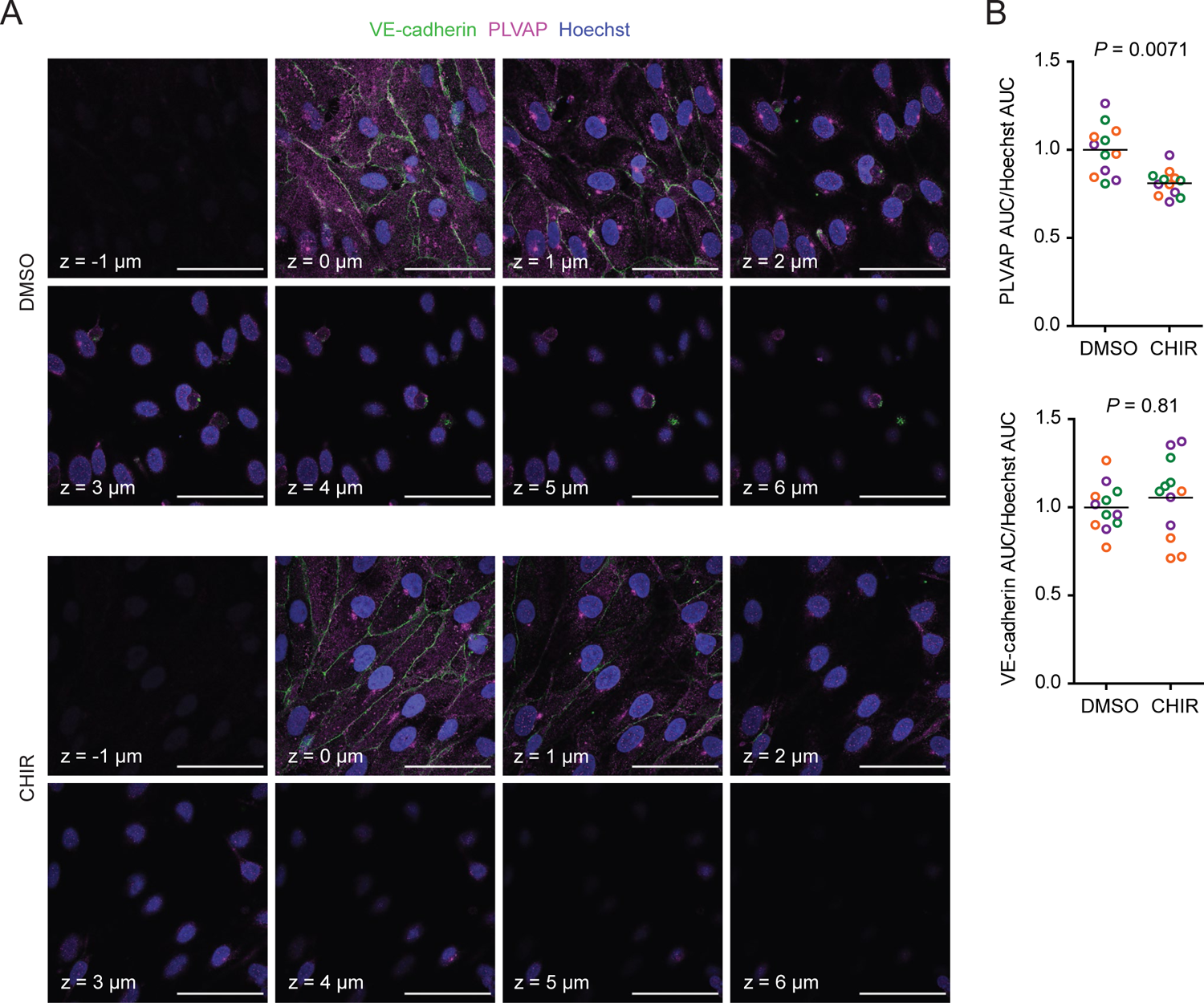
Effect of CHIR on endothelial PLVAP expression. **(A)** Confocal immunocytochemistry analysis of VE-cadherin and PLVAP expression in Passage 1 ECs treated with DMSO or CHIR. Hoechst nuclear counterstain is overlaid. Eight serial confocal Z-slices with 1 μm spacing are shown. Scale bars: 50 μm. **(B)** Quantification of PLVAP and VE-cadherin area under the curve (AUC) of mean fluorescence intensity versus Z-position normalized to Hoechst AUC. Points represent replicate wells from 3 independent differentiations of the IMR90-4 line, each differentiation indicated with a different color. Bars indicate mean values, with values normalized within each differentiation such that the mean of the DMSO condition equals 1. P-values: Two-way ANOVA on unnormalized data.

**Figure 5.**
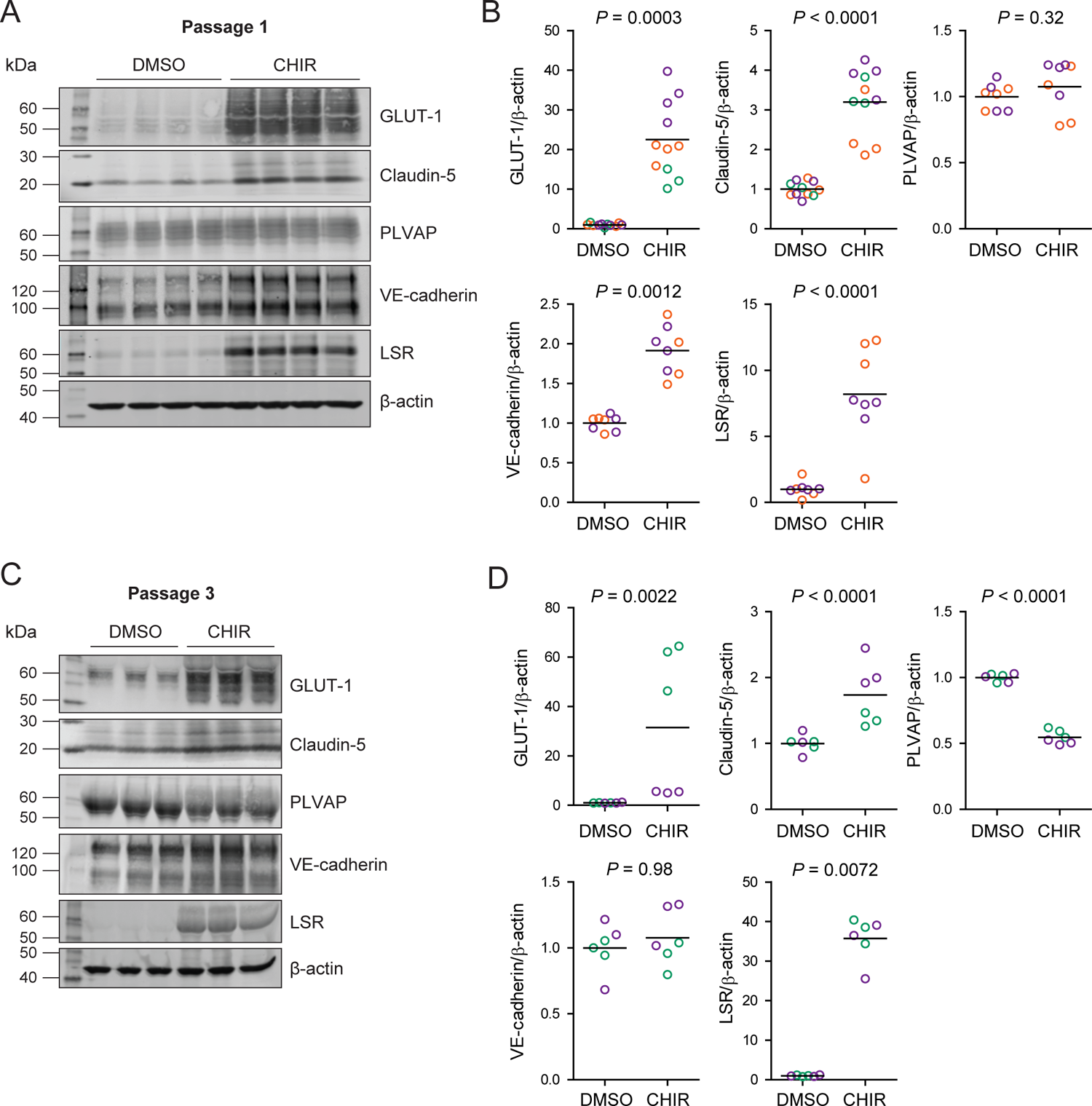
Effect of CHIR on protein expression in Passage 1 and Passage 3 ECs. **(A)** Western blots of Passage 1 ECs treated with DMSO or CHIR probed for GLUT-1, claudin-5, PLVAP, VE-cadherin, LSR, and β-actin. **(B)** Quantification of Western blots of Passage 1 ECs. GLUT-1, claudin-5, PLVAP, VE-cadherin, and LSR band intensities were normalized to β-actin band intensity. Points represent replicate wells from 2–3 independent differentiations of the IMR90-4 line, each differentiation indicated with a different color. Bars indicate mean values, with values normalized within each differentiation such that the mean of the DMSO condition equals 1. P-values: Two-way ANOVA on unnormalized data. **(C)** Western blots of Passage 3 ECs treated with DMSO or CHIR probed for GLUT-1, claudin-5, PLVAP, VE-cadherin, LSR, and β-actin. **(D)** Quantification of Western blots of passage 3 ECs. GLUT-1, claudin-5, PLVAP, VE-cadherin, and LSR band intensities were normalized to β-actin band intensity. Points represent replicate wells from 2 independent differentiations of the IMR90-4 line, each differentiation indicated with a different color. Bars indicate mean values, with values normalized within each differentiation such that the mean of the DMSO condition equals 1. P-values: Two-way ANOVA on unnormalized data.

CHIR treatment produced two apparently competing changes in ECs related to vesicular transport: an expected downregulation of PLVAP and an unexpected upregulation of caveolin-1. We therefore asked whether the rate of total fluid-phase endocytosis differed between CHIR-treated and control ECs, using a fluorescently-labeled 10 kDa dextran as a tracer. After incubating Passage 1 cultures with dextran for 2 h at 37°C, we used flow cytometry to gate CD31^+^ ECs and assess total dextran accumulation (Figure 6A-B). We first confirmed that the process of dextran internalization required the membrane fluidity of an endocytosis-dependent process by carrying out the assay at 4°C; this condition indeed yielded a substantially decreased dextran signal compared to 37°C (Figure 6B). In ECs incubated at 37°C, CHIR treatment did not change the geometric mean dextran signal compared to DMSO (Figure 6B,C), but did cause a broadening of the distribution of dextran intensities, indicative of sub-populations of cells with decreased and increased dextran uptake (Figure 6B,D). Importantly, these results were consistent across three independent differentiations (Figure 6C-D). Thus, despite the generally uniform elevation of caveolin-1 and decrease of PLVAP observed by immunocytochemistry in CHIR-treated ECs, our functional assay suggests neither an overall increase nor decrease in total fluid-phase endocytosis. Instead, it indicates that CHIR increases the heterogeneity of the EC population with respect to the rate of endocytosis.

**Figure 6.**
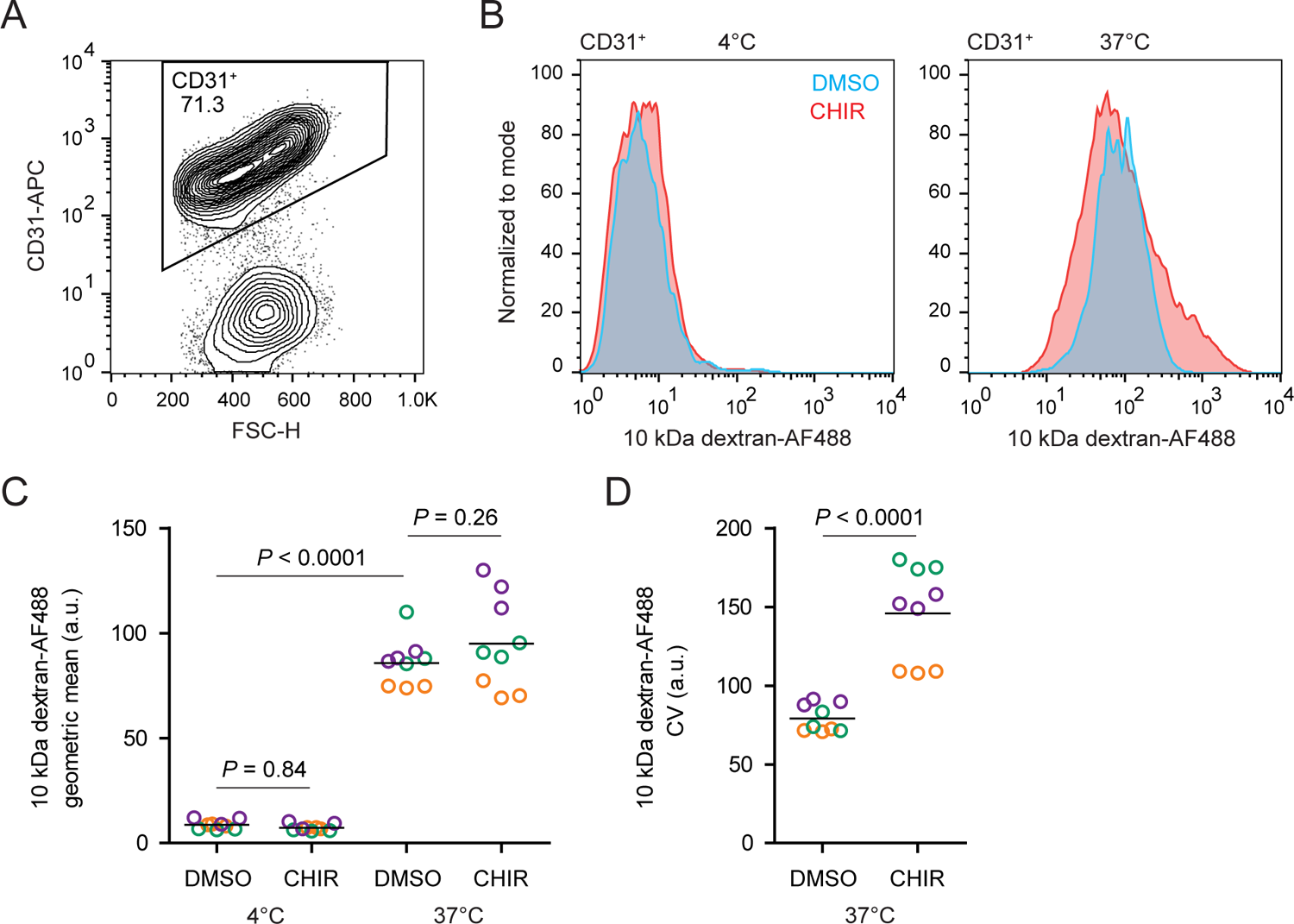
Fluid-phase endocytosis in CHIR- and DMSO-treated ECs. **(A)** Flow cytometry analysis of CD31 expression in Passage 1 ECs following the dextran internalization assay. CD31^+^ cells were gated for further analysis. **(B)** Flow cytometry analysis of 10 kDa dextran-Alexa Fluor 488 (AF488) abundance in CD31^+^ cells. Cells were treated with DMSO or CHIR for 6 d prior to the assay. Representative plots from cells incubated with dextran for 2 h at 4°C (left) and 37°C (right) are shown. **(C)** Quantification of 10 kDa dextran-AF488 geometric mean fluorescence intensity in CD31^+^ cells. Treatment and assay conditions were as described in (B). Points represent replicate wells from 3 independent differentiations of the IMR90-4 line, each differentiation indicated with a different color. Bars indicate mean values. P-values: Two-way ANOVA followed by Tukey’s HSD test. **(D)** Quantification of the coefficient of variation (CV) of 10 kDa dextran-AF488 fluorescence intensity in CD31^+^ cells. Points represent replicate wells from 3 independent differentiations of the IMR90-4 line, each differentiation indicated with a different color. Bars indicate mean values. P-value: Two-way ANOVA.

Given the relatively weak responses to Wnt activation in adult mouse liver ECs *in vivo* (Munji et al., 2019) and adult mouse brain ECs cultured *in vitro* (Sabbagh and Nathans, 2020), we sought to determine whether the immature, potentially more plastic state of hPSC-derived endothelial progenitors contributed to the relatively robust CHIR-mediated response we observed. To test this hypothesis, we matured hPSC-derived ECs *in vitro* for 4 passages (until approximately day 30) prior to initiating CHIR treatment for 6 days (Figure 7A). The resulting Passage 5 DMSO-treated ECs, which are analogous to EECM-BMEC-like cells we previously reported (Nishihara et al., 2020), did not have detectable GLUT-1 expression (Figure 7B). Compared to DMSO controls, the resulting CHIR-treated Passage 5 ECs exhibited an approximately 1.5-fold increase in GLUT-1 abundance (Figure 7B-C), a markedly weaker response than the 10- to 40-fold increases routinely observed using the same immunocytochemistry-based assay when CHIR treatment was initiated immediately after MACS (Figure 2; Figure 2–figure supplement 1; Figure 2–figure supplement 2; Figure 3). Furthermore, CHIR treatment in matured ECs led to a slight decrease in EC number (Figure 7D), rather than the increase observed when treatment was initiated immediately after MACS (Figure 2–figure supplement 1; Figure 3). Together, these data suggest that early, naïve endothelial progenitors are more responsive to Wnt activation than more mature ECs derived by the same differentiation protocol.

**Figure 7.**
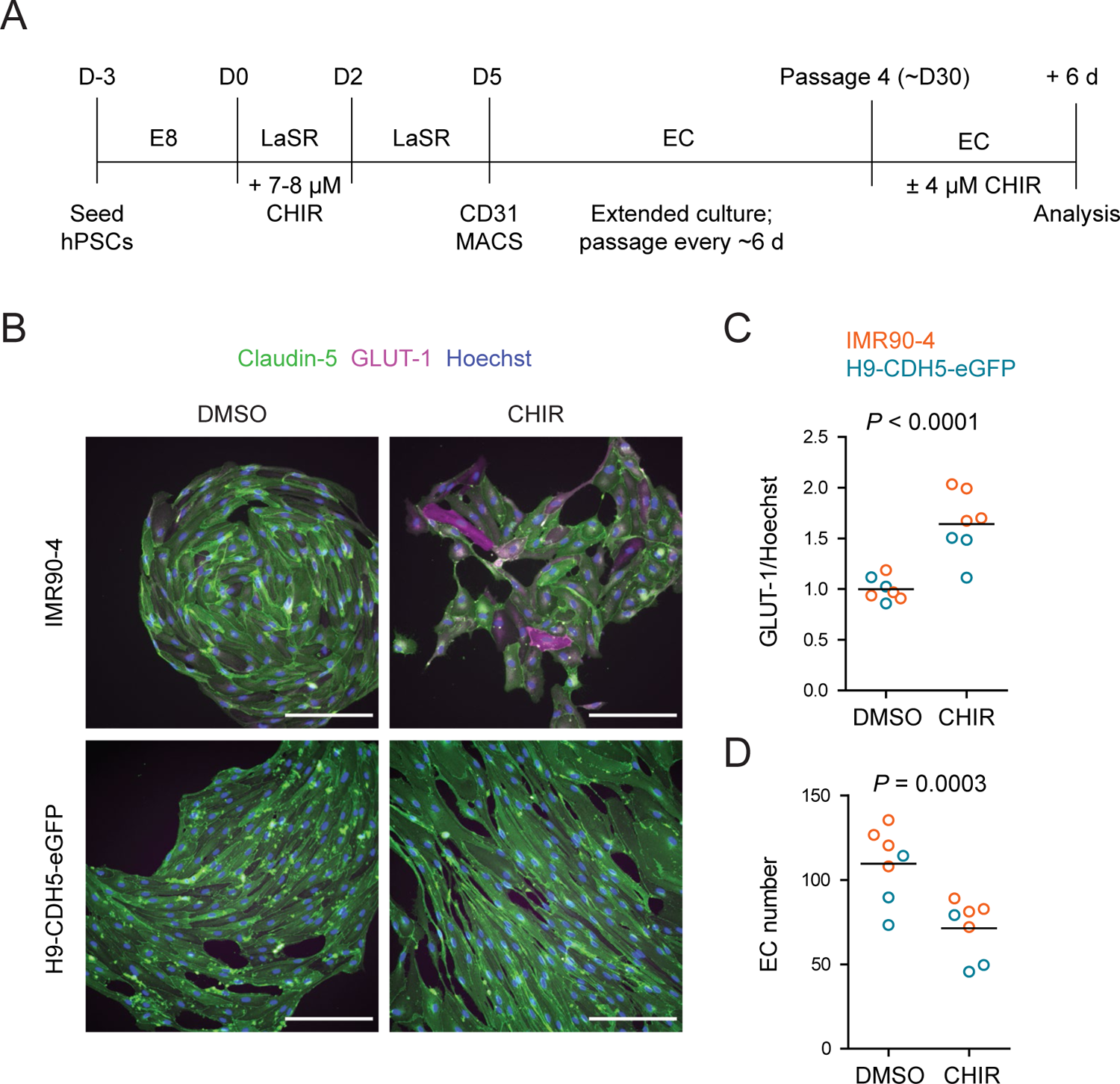
Effect of CHIR treatment in matured endothelium. (**A**) Overview of the endothelial differentiation, extended culture, and CHIR treatment protocol. **(B)** Immunocytochemistry analysis of claudin-5 and GLUT-1 expression in ECs treated with DMSO or CHIR as outlined in (A). Images from the IMR90-4 and H9-CDH5-eGFP lines are shown. Hoechst nuclear counterstain is overlaid. Scale bars: 200 μm. **(C)** Quantification of images from the conditions described in (B) for GLUT-1 mean fluorescence intensity normalized to Hoechst mean fluorescence intensity within the area of claudin-5^+^ ECs only. Points represent replicate wells from one differentiation of the IMR90-4 line (orange) and one differentiation of the H9-CDH5-eGFP line (blue). Bars indicate mean values, with values normalized within each differentiation such that the mean of the DMSO condition equals 1. P-value: Two-way ANOVA on unnormalized data. **(D)** Quantification of images from the conditions described in (B) for number of ECs per 20× field. Points represent replicate wells from one differentiation of the IMR90-4 line (orange) and one differentiation of the H9-CDH5-eGFP line (blue). Bars indicate mean values. P-value: Two-way ANOVA.

### Comprehensive profiling of the Wnt-regulated endothelial transcriptome

We turned next to RNA-sequencing as an unbiased method to assess the impacts of Wnt activation on the EC transcriptome. We performed four independent differentiations and analyzed Passage 1 ECs treated with DMSO, CHIR, or Wnt7a and Wnt7b (Wnt7a/b), using fluorescence-activated cell sorting (FACS) to isolate CD31^+^ ECs from the mixed EC/SMLC cultures. We also sequenced the SMLCs from DMSO-treated cultures at Passage 1 from two of these differentiations. DMSO- and CHIR-treated ECs at Passage 3 from three of these differentiations were also sequenced. Principal component analysis of the resulting whole-transcriptome profiles revealed that the two cell types (ECs and SMLCs) segregated along principal component (PC) 1, which explained 52% of the variance. In ECs, the effects of passage number and treatment were reflected in PC 2, which explained 20% of the variance (Figure 8A). We next validated the endothelial identity of our cells; we observed that canonical endothelial marker genes (including *CDH5, CD34, PECAM1, CLDN5, ERG,* and *FLI1*) were enriched in ECs compared to SMLCs and had high absolute abundance, on the order of 100–1,000 transcripts per million (TPM) (Figure 8B; Supplementary file 1). SMLCs expressed mesenchymal (mural/fibroblast)-related transcripts (including *PDGFRB, CSPG4, PDGFRA, TBX2, CNN1,* and *COL1A1*), which ECs generally lacked, although we did observe slight enrichment of some of these genes in Passage 1 DMSO-treated ECs, likely reflective of a small amount of SMLC contamination despite CD31 FACS (Figure 8B). SMLCs also expressed *SLC2A1* (Supplementary file 1) consistent with protein-level observations (Figure 1D). We also observed little to no expression of the epithelial genes *CDH1*, *EPCAM*, *CLDN1*, *CLDN3* (Castro Dias et al., 2019), *CLDN4*, and *CLDN6*, reflecting the definitive endothelial nature of the cells (Figure 8B; Supplementary file 1).

**Figure 8.**
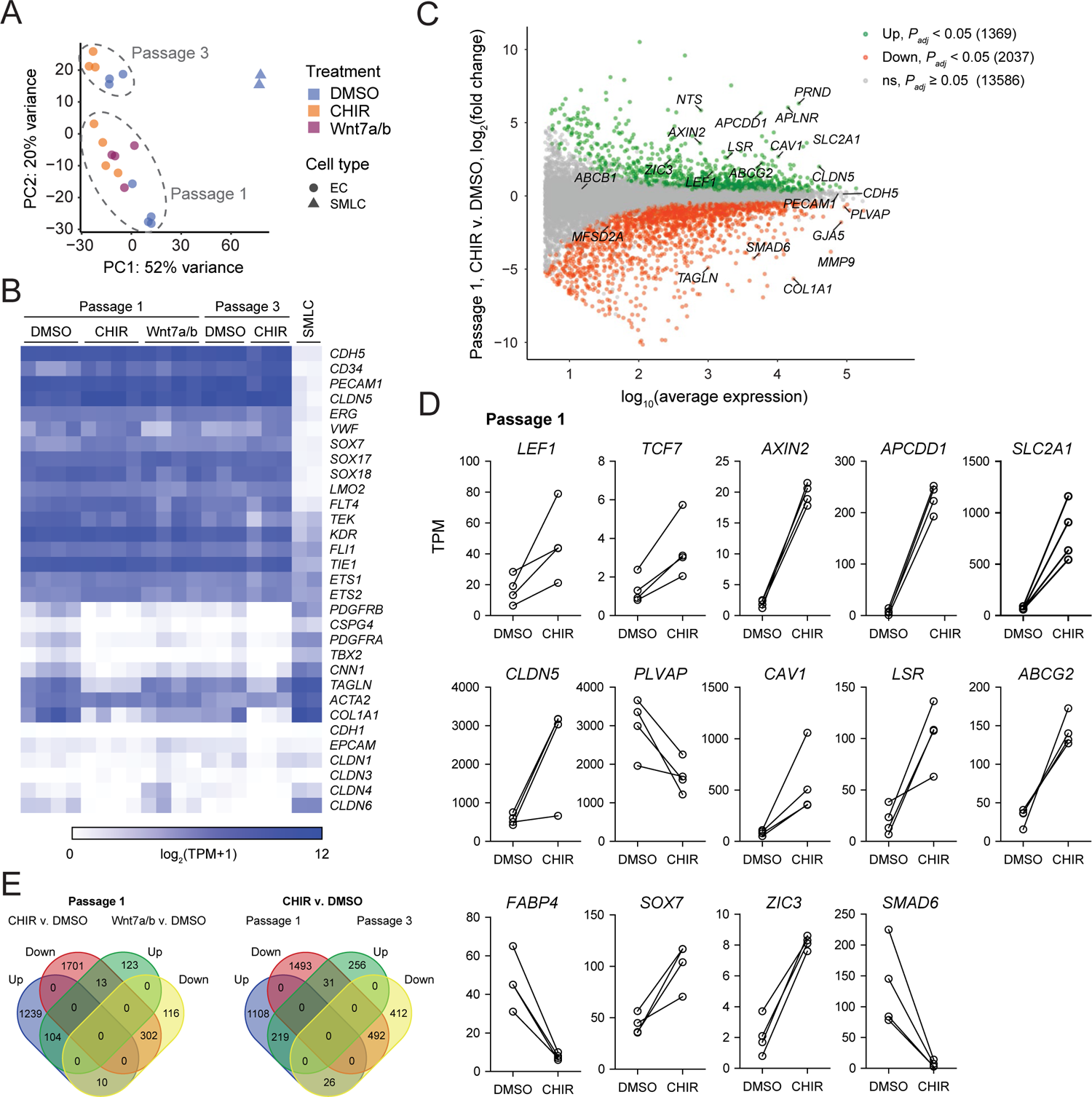
RNA-seq of DMSO-, CHIR-, or Wnt7a/b-treated ECs. **(A)** Principal component analysis of EC and SMLC whole-transcriptome data subject to variance stabilizing transformation by DESeq2. Points from Passage 1 ECs represent cells from 4 independent differentiations of the IMR90-4 line, points from Passage 3 ECs represent cells from 3 independent differentiations of the IMR90-4 line, and points from SMLCs represent 2 independent differentiations of the IMR90-4 line. Points are colored based on treatment: DMSO (blue), CHIR (orange), or Wnt7a/b (red). Data are plotted in the space of the first two principal components, with the percentage of variance explained by principal component 1 (PC1) and principal component 2 (PC2) shown in axis labels. **(B)** Heat map of transcript abundance [log_2_(TPM+1)] for endothelial, mesenchymal, and epithelial genes across all samples. Abundance data for all transcripts is provided in Supplementary file 1. **(C)** Differential expression analysis of Passage 1 CHIR-treated ECs compared to Passage 1 DMSO-treated ECs. Differentially expressed genes (adjusted P-values < 0.05, DESeq2 Wald test with Benjamini-Hochberg correction) are highlighted in green (upregulated) and red (downregulated). The number of upregulated, downregulated, and non-significant (ns) genes are shown in the legend. Complete results of differential expression analysis are provided in Supplementary file 2. **(D)** Transcript abundance (TPM) of Wnt-regulated, barrier-related genes in Passage 1 DMSO- and CHIR-treated ECs. Points represent cells from 4 independent differentiations of the IMR90-4 line and lines connect points from matched differentiations. All genes shown were differentially expressed (adjusted P-values < 0.05, DESeq2 Wald test with Benjamini-Hochberg correction). P-values are provided in Supplementary file 2. **(E)** Venn diagrams illustrating the number of genes identified as upregulated or downregulated (adjusted P-values < 0.05, DESeq2 Wald test with Benjamini-Hochberg correction) in Passage 1 ECs treated with CHIR versus DMSO compared to Wnt7a/b versus DMSO (left), or ECs treated with CHIR versus DMSO at Passage 1 compared to Passage 3 (right). Gene lists are provided in Supplementary file 2.

**Figure 8–figure supplement 1.**
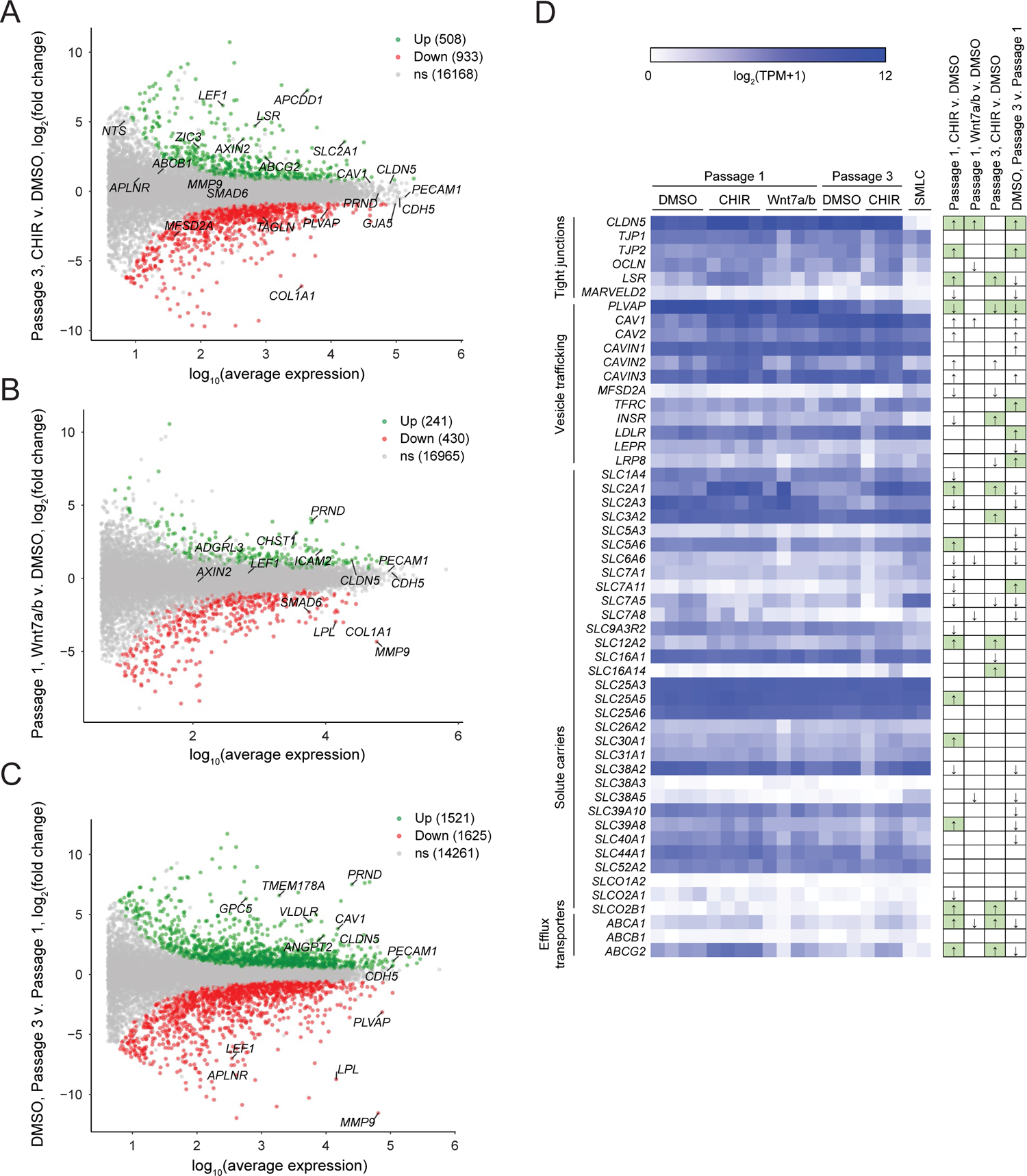
RNA-seq differential expression analyses. **(A-C)** Differential expression analysis of Passage 3 CHIR-treated ECs compared to Passage 3 DMSO-treated ECs (A), Passage 1 Wnt7a/b-treated ECs compared to Passage 1 DMSO-treated ECs (B), and Passage 3 DMSO-treated ECs compared to Passage 1 DMSO-treated ECs (C). Differentially expressed genes (adjusted P-values < 0.05, DESeq2 Wald test with Benjamini-Hochberg correction) are highlighted in green (upregulated) and red (downregulated). The number of upregulated, downregulated, and non-significant (ns) genes are shown in the legends. Complete results of differential expression analyses are provided in Supplementary file 2. **(D)** Heat map of transcript abundance [log_2_(TPM+1)] for BBB genes encompassing tight junctions, vesicle trafficking components, and solute carriers, and efflux transporters. Solute carrier and efflux transporter genes that were expressed in human brain ECs at an average of >100 TPM in a meta-analysis of scRNA-seq datasets (Gastfriend et al., 2021) are included. Abundance data for all transcripts is provided in Supplementary file 1. At right, arrows indicate directionality of change for differentially expressed genes (adjusted P-values < 0.05, DESeq2 Wald test with Benjamini-Hochberg correction) for the four comparisons shown above. Changes with expected directionality for gain of CNS EC character have arrows highlighted in green.

**Figure 8–figure supplement 2.**
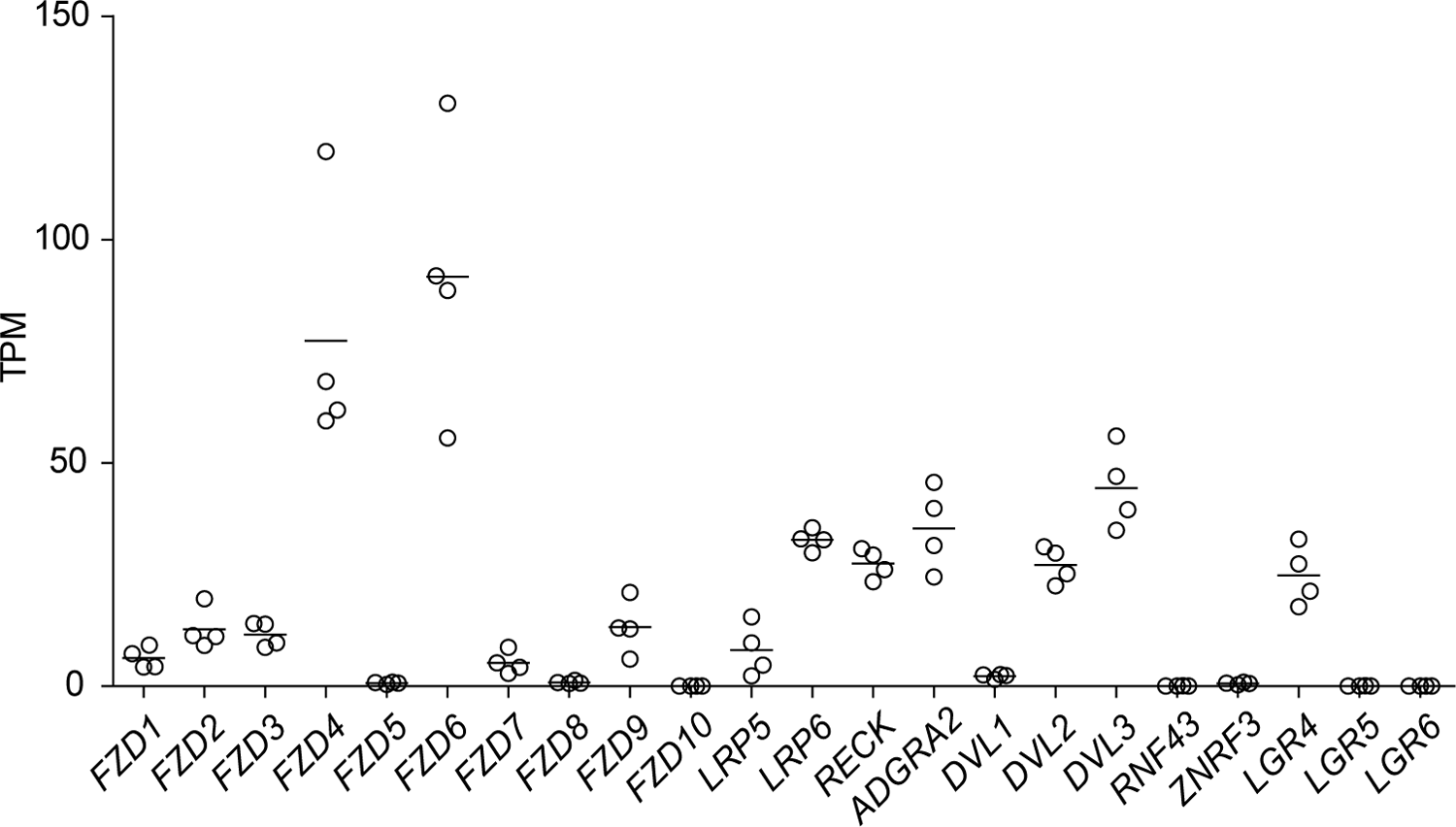
Expression of Wnt pathway components in naïve ECs. Abundance of transcripts (in transcripts per million, TPM) encoding Wnt receptors, co-receptors, and other pathway components in Passage 1 DMSO-treated ECs. Points represent cells from 4 independent differentiations of the IMR90-4 line. Bars indicate mean values. *ADGRA2* is also known as *GPR124*.

First comparing CHIR- and DMSO-treated ECs at Passage 1, we identified 1,369 significantly upregulated genes and 2,037 significantly downregulated genes (Figure 8C; Supplementary file 2). CHIR-upregulated genes included *SLC2A1*, *CLDN5*, *LSR*, and *CAV1*, consistent with protein-level assays. *PLVAP* was downregulated, as were a number of mesenchymal genes (*TAGLN*, *COL1A1*), again reflective of slight contamination of SMLC transcripts in the DMSO-treated EC samples (Figure 8C-D). Additionally, important downstream effectors of Wnt signaling were upregulated, including the transcription factors *LEF1* and *TCF7*, the negative regulator *AXIN2*, and the negative regulator *APCDD1*, which is known to modulate Wnt-regulated barriergenesis in retinal endothelium (Mazzoni et al., 2017) (Figure 8C-D). We also found that the transcription factors *ZIC3,* which is highly enriched in brain and retinal ECs *in vivo* and downstream of Frizzled4 signaling (Wang et al., 2012; Sabbagh et al., 2018), and *SOX7*, which acts cooperatively with *SOX17* and *SOX18* in retinal angiogenesis (Zhou et al., 2015), were upregulated by CHIR in our system (Figure 8D). Additional CHIR-upregulated genes included *ABCG2* (encoding the efflux transporter Breast Cancer Resistance Protein, BCRP), and *APLN*, a tip cell marker enriched in postnatal day 7 murine brain ECs compared to those of other organs, and subsequently downregulated in adulthood (Sabbagh et al., 2018; Sabbagh and Nathans, 2020) (Figure 8C). Finally, we detected CHIR-mediated downregulation of the fatty acid-binding protein-encoding *FABP4*, which is depleted in brain ECs compared to those of peripheral organs (Sabbagh et al., 2018). We also observed similar downregulation of *SMAD6*, which is depleted in brain ECs compared to lung ECs and is a putative negative regulator of BMP-mediated angiogenesis (Mouillesseaux et al., 2016; Vanlandewijck et al., 2018) (Figure 8D). Many of these CHIR-mediated gene expression changes persisted at Passage 3, including *SLC2A1*, *LSR*, *LEF1*, *AXIN2*, *APCDD1*, *ZIC3*, and *ABCG2* upregulation and *PLVAP* downregulation (Figure 8E; Figure 8–figure supplement 1A).

We made similar comparisons (i) between Wnt7a/b-treated and control (DMSO-treated) ECs at Passage 1, and (ii) between control ECs at Passage 3 versus Passage 1 (Figure 8E; Figure 8–figure supplement 1B-C; Supplementary file 2). Consistent with the weak response observed by immunocytochemistry, there were fewer Wnt7a/b-mediated gene expression changes compared to those elicited by CHIR, with 241 upregulated and 420 downregulated genes (Figure 8–figure supplement 1B). In general, however, these changes were consistent with CHIR-mediated changes, with 104 concordantly upregulated genes, 302 concordantly downregulated genes, and only 23 discordantly regulated genes (Figure 8E). Of note, treatment with Wnt7a/b, but not CHIR, upregulated *SOX17*, a Wnt target gene required for BBB function (Corada et al., 2018). Extended culture to Passage 3 in the absence of exogeneous Wnt activation led to 1,521 upregulated genes, including *CLDN5* and *CAV1*, consistent with previously-reported protein-level observations in EECM-BMEC-like cells (Nishihara et al., 2020), which are analogous to Passage 3 DMSO-treated cells. We also observed 1,625 downregulated genes, including *PLVAP* (Figure 8–figure supplement 1C). *SLC2A1,* however, was not upregulated at Passage 3 (Figure 8–figure supplement 1C), concordant with absence of GLUT-1 protein expression in the control ECs (Figure 7B). To further understand the strengths and limitations of this model system both as a readout of early developmental changes in CNS ECs (Passage 1 cells) or as a source of CNS-like ECs for use in downstream modeling applications, we evaluated absolute transcript abundance and effects of treatment or passage number on 53 characteristic CNS EC genes encompassing tight junction components, vesicle trafficking machinery, solute carriers, and ATP-binding cassette (ABC) efflux transporters selected based on high expression in human brain endothelial cells from a meta-analysis of single cell RNA-seq data (Gastfriend et al., 2021) (Figure 8–figure supplement 1D). While ECs expressed *CLDN5*, *TJP1*, *TJP2*, *OLCN*, and *LSR*, they lacked *MARVELD2* (encoding tricellulin) under all conditions. ECs under all conditions also lacked *MFSD2A* and, despite CHIR-mediated downregulation of *PLVAP*, retained high absolute expression of this and other caveolae-associated genes. Finally, while many solute carriers and ABC transporters were expressed (*SLC2A1*, *SLC3A2*, *SLC16A1*, *SLC38A2*, *ABCG2*), others expressed at the *in vivo* human BBB were not (*SLC5A3*, *SLC7A11*, *SLC38A3*, *SLCO1A2*, *ABCB1*) (Figure 8–figure supplement 1D). Thus, while CHIR treatment yields ECs with certain elements of CNS-like character, additional molecular signals are likely necessary to improve other aspects of the *in vivo* CNS EC phenotype.

To partially address the hypothesis that the weak response of ECs to Wnt7a/b, NR-CM, and Astro-CM is due to a lack of necessary Wnt receptors and/or co-receptors, we used RNA-seq data from Passage 1 DMSO-treated ECs to evaluate expression of transcripts encoding these and other components of the canonical Wnt signaling pathway. *FZD4* and *FZD6* were highly expressed and enriched compared to all other Frizzleds (Figure 8–figure supplement 2), consistent with data from murine brain ECs *in vivo* (Daneman et al., 2009). *RECK* and *ADGRA2* (*GPR124*) were moderately expressed at a level similar to *LRP6* (on the order of 40 TPM), while little to no *LRP5* was expressed (Figure 8–figure supplement 2). Taken together, however, these data suggest that the hPSC-derived ECs express much of the machinery necessary to transduce the signal from Wnt7a/b ligands, but possibilities remain that the proteins encoded by the evaluated transcripts are absent, or at too low an abundance, for a robust response, motivating the use of CHIR to bypass the cell surface Wnt pathway components for robust induction of barriergenesis via β-catenin stabilization.

### The Wnt-regulated endothelial transcriptome in multiple contexts

To globally assess whether CHIR-mediated gene expression changes in our system are characteristic of the responses observed in ECs *in vivo* and similar to those observed in other *in vitro* contexts, we compared our RNA-seq dataset to those of studies that employed a genetic strategy for β-catenin stabilization (the *Ctnnb1*^flex3^ allele) in adult mouse ECs in several contexts: (i) pituitary ECs, which acquire some BBB-like properties upon β-catenin stabilization (Wang et al., 2019); (ii) liver ECs, which exhibit little to no barriergenic response to β-catenin stabilization (Munji et al., 2019); (iii) brain ECs briefly cultured *in vitro*, which rapidly lose their BBB-specific gene expression profile even with β-catenin stabilization (Sabbagh and Nathans, 2020), and offer the most direct comparison to our *in vitro* model system. Upon recombination, the *Ctnnb1*^flex3^ allele produces a dominant mutant β-catenin lacking residues that are phosphorylated by GSK-3β to target β-catenin for degradation (Harada et al., 1999); as such, this strategy for ligand- and receptor-independent Wnt activation by β-catenin stabilization is directly analogous to CHIR treatment.

We first used literature RNA-seq data from postnatal day 7 murine brain, liver, lung, and kidney ECs (Sabbagh et al., 2018) to define core sets of genes in brain ECs that are differentially expressed compared to all three of the other organs (Figure 9A-B). Using the resulting sets of 1094 brain-enriched and 506 brain-depleted genes, we asked how many genes in our Passage 1 ECs were concordantly-regulated by CHIR: 130 of the brain-enriched genes were CHIR-upregulated and 116 of the brain-depleted genes were CHIR-downregulated (Figure 9C). In pituitary ECs with β-catenin stabilization, 102 of the brain enriched genes were upregulated with and 48 of the brain depleted genes were downregulated (Figure 9D). Compared with the pituitary ECs, there were far fewer concordantly-regulated genes in liver ECs with β-catenin stabilization, with 25 upregulated and 1 downregulated (Figure 9E). Finally, cultured primary mouse brain ECs with β-catenin stabilization exhibited 72 upregulated and 16 downregulated genes (Figure 9F). The only gene concordantly-regulated in all four comparisons was the canonical Wnt target *AXIN2*. Several additional genes were concordantly upregulated in three of four, including *TCF7*, *FAM107A*, *NKD1*, *TNFRSF19*, *GLUL*, *SLC30A1*, and *ABCB1*, which was the only gene concordantly regulated in all comparisons except the hPSC-derived ECs (Figure 9G). Several canonical target genes were shared by the hPSC-derived EC and pituitary EC systems, including *APCDD1*, *LEF1*, *CLDN5*, and *SLC2A1*; also in this category were *LSR*, the zinc/manganese transporter *SLC39A8*, and 12 additional genes (Figure 9G). Notably, the caveolae inhibitor *MFSD2A* was robustly upregulated by β-catenin in pituitary ECs, but not in any other context (Figure 9C-F), suggesting other brain-derived factors may cooperate with Wnt to regulate expression of this important inhibitor of caveolin-mediated transcytosis. Complete gene lists from this comparative analysis are provided in Supplementary file 3. In sum, the data suggest that the hPSC-derived ECs responded to Wnt activation in a fashion that led to modest induction of CNS transcriptional programs and that the response was most similar to the pituitary β-catenin stabilization model. Importantly, this analysis also supports the hypothesis that immature endothelium is highly responsive to Wnt activation where mature (adult) endothelium is largely refractory except in regions proximal to barrier-forming regions.

**Figure 9.**
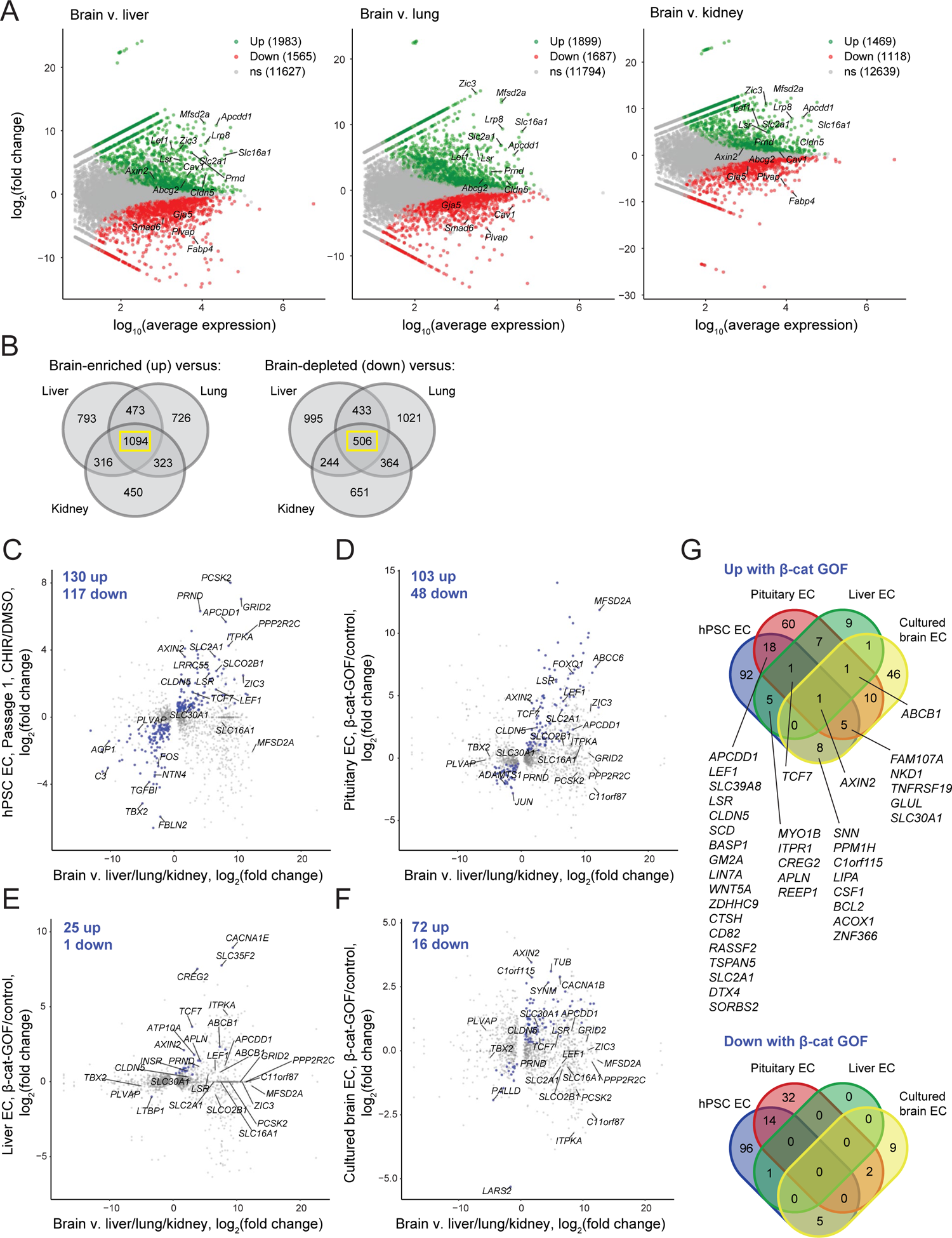
Identification of concordantly Wnt-regulated CNS EC-associated genes in RNA-seq data. **(A)** Differential expression analysis of P7 murine brain ECs compared to liver, lung, or kidney ECs (Sabbagh *et al*., 2018). Differentially expressed genes (adjusted P-values < 0.05, DESeq2 Wald test with Benjamini-Hochberg correction) are highlighted in green (up, brain-enriched) and red (down, brain-depleted). The number of up, down, and non-significant (ns) genes are shown in the legends. **(B)** Venn diagrams illustrating the number of genes identified as brain EC-enriched (left) or brain EC-depleted (right) versus liver, lung, or kidney ECs (adjusted P-values < 0.05, DESeq2 Wald test with Benjamini-Hochberg correction). The 1094 genes enriched in brain ECs compared to each other organ, and the 506 genes depleted in brain ECs compared to each other organ, were used for subsequent analysis of the effects of Wnt activation in the various experimental contexts. **(C-F)** In each plot, the x-axis indicates average log_2_(fold change) of gene expression in brain ECs compared to liver, lung, and kidney ECs for the 1094 brain EC-enriched genes and 506 brain EC-depleted genes described in with known mouse-human homology. Homologous human gene names are shown. The y-axes indicate differential expression [log_2_(fold change)] in Passage 1 CHIR-treated ECs compared to Passage 1 DMSO-treated ECs (C), in adult mouse pituitary ECs with stabilized β-catenin (gain-of-function, GOF) compared to controls (Wang *et al*., 2019) (D), in adult mouse liver ECs with stabilized β-catenin compared to controls (Munji *et al*., 2020) (E), or in cultured adult mouse brain ECs with stabilized β-catenin compared to controls (Sabbagh *et al*., 2020) (F). Points are highlighted in blue if concordantly-regulated (upregulated in both comparisons or downregulated in both comparisons). Genes were identified as upregulated or downregulated based on adjusted P-values < 0.05, DESeq2 Wald test with Benjamini-Hochberg correction. **(G)** Venn diagrams illustrating the number of brain EC-enriched genes concordantly upregulated with β-catenin GOF (top) and the number of brain EC-depleted genes concordantly downregulated with β-catenin GOF (bottom) for the four comparisons shown in (C-F). Complete results of this analysis are provided in Supplementary file 3.

## DISCUSSION

The Wnt/β-catenin signaling pathway plays a central role in CNS angiogenesis and in establishing the unique properties of CNS ECs (Liebner et al., 2008; Stenman et al., 2008; Daneman et al., 2009; Kuhnert et al., 2010; Cullen et al., 2011; Vanhollebeke et al., 2015; Cho et al., 2017). In this work, we investigated the role of Wnt/β-catenin signaling on induction of BBB properties in a human EC model, using naïve endothelial progenitors derived from hPSCs. We reasoned that these immature EPCs (Lian et al., 2014) would be similar to the immature endothelium in the perineural vascular plexus and thus competent to acquire CNS EC phenotypes in response to Wnt activation. We evaluated several strategies to activate Wnt, including the widely used ligand Wnt3a (Liebner et al., 2008), the neural progenitor- and astrocyte-derived ligands Wnt7a and Wnt7b, which are the two Wnt ligands primarily responsible for the Wnt-dependent effects of CNS angiogenesis and barriergenesis observed *in vivo* (Daneman et al., 2009; Cho et al., 2017), neural rosette- and astrocyte-CM as putative cellular sources of Wnt ligands, and the GSK-3 inhibitor CHIR.

We found that CHIR treatment robustly induced several canonical CNS EC molecular phenotypes, including a marked induction of GLUT-1, upregulation of claudin-5, and downregulation of PLVAP, which correlated with differential gene expression in RNA-seq data. Further, using RNA-seq and Western blotting, we also identified LSR (angulin-1) as CHIR-induced in this system, supporting the notion that this highly CNS EC-enriched tricellular tight junction protein (Daneman et al., 2010a; Sohet et al., 2015) is Wnt-regulated. In RNA-seq data, we observed differential expression of known CNS EC-enriched/depleted and Wnt-regulated genes including upregulated *LEF1*, *AXIN2*, *APCDD1*, *ABCG2*, *SOX7*, and *ZIC3* and downregulated *PLVAP*, *FABP4*, and *SMAD6*. These RNA-seq data should therefore be useful in generating hypotheses of BBB-associated genes regulated by Wnt activation in ECs, for future functional studies. Our work also defines an important set of phenotypes for which Wnt activation in ECs is not sufficient in our system: in the context of vesicle trafficking, we observed caveolin-1 (*CAV1*) upregulation, no change in mean functional endocytosis, virtually no expression of *MFSD2A*, and high absolute *PLVAP* abundance despite CHIR-mediated downregulation. Given roles of brain pericytes in regulating PLVAP, MFSD2A, and functional transcytosis (Armulik et al., 2010; Daneman et al., 2010b; Ben-Zvi et al., 2014; Stebbins et al., 2019), and the observation that MFSD2A is Wnt-regulated in pituitary ECs *in vivo* (Wang et al., 2019), where pericytes are present, it is plausible that pericyte-derived cues are necessary in addition to Wnts to achieve the characteristically low rate of CNS EC pinocytosis. Next, while *ABCG2* (BCRP) was Wnt-induced in our system, other hallmark efflux transporters were not Wnt-regulated and either expressed at low levels (e.g., *ABCC4*, encoding MRP-4) or not expressed (e.g. *ABCB1*, encoding P-glycoprotein). Notably however, *Abcb1a* was Wnt-regulated in the three other β-catenin stabilization experiments from the literature that we evaluated (Munji et al., 2019; Wang et al., 2019; Sabbagh and Nathans, 2020). Thus, pericyte-derived cues, astrocyte-derived cues, and/or activation of the pregnane X or other nuclear receptors may be important for complete acquisition of the complement of CNS EC efflux transporters (Bauer et al., 2004; Berezowski et al., 2004; Praça et al., 2019).

While several recombinant Wnt ligands and neural rosette-CM elevated GLUT-1 expression in ECs, the magnitude of this effect was small compared to the robust induction of GLUT-1 observed with CHIR treatment. While we observed moderate transcript-level expression *RECK* and *ADGRA2* (*GPR124*) in Passage 1 ECs, it is possible that protein-level expression of these necessary Wnt7 coreceptors, or additional components necessary for Wnt signal transduction, are not of sufficient abundance. For example, absence of LRP5 is a potential factor in the muted response to Wnt ligands and CM because LRP5 and LRP6 likely have non-redundant functions, as evidenced by defects in retinal barrier formation in *Lrp5-*knockout mice (Zhou et al., 2014). Presence of GPR124 in naïve endothelial progenitors is consistent with ubiquitous expression in ECs in the mouse embryo that is subsequently downregulated in non-CNS endothelium; however, GPR124 enrichment in CNS ECs can be observed as early as E12.5 (Kuhnert et al., 2010), leaving open the possibility that during development other neural tissue-derived signals upregulate or maintain RECK and GPR124 expression. Furthermore, while ligand potency or concentration may also play a role in the weak response, we observed a consistent and highly potent EC-purifying effect (i.e., reduction or elimination of the contaminating SMLCs observed in control Passage 1 cultures) with Wnt7a and both neural rosette- and astrocyte-CM. CHIR also achieved this purifying effect and increased EC number, suggesting that Wnt signaling plays a role in suppressing proliferation of mesoderm-derived mural cells in this system.

We also directly addressed the hypothesis that immature ECs are more plastic, that is, more competent to acquire BBB properties upon Wnt activation, than mature ECs. This hypothesis is supported by existing observations that ectopic expression of Wnt7a is sufficient to induce GLUT-1 expression in non-CNS regions of the mouse embryo (Stenman et al., 2008), but β-catenin stabilization in adult mouse liver and lung ECs produces only a slight effect (Munji et al., 2019). We repeated our CHIR treatment paradigm in hPSC-derived ECs after an extended period of *in vitro* culture, and observed much weaker induction of GLUT-1 and no pro-proliferative effect. Thus, our results support this hypothesis and suggest that the loss of BBB developmental plasticity in ECs is an intrinsic, temporally-controlled process rather than a result of the peripheral organ environment. Interestingly, ECs in non-BBB-forming regions of the CNS (i.e., CVOs), and in the anterior pituitary, which is directly proximal to the CNS, retain some of their plasticity in adulthood (Wang et al., 2019), possibly as the result of a delicate balance between Wnt ligands and Wnt-inhibitory factors in these regions. Our model should facilitate additional systematic examination of factors that may enhance or attenuate EC Wnt responsiveness.

Finally, our work establishes an improved hPSC-based model for investigating mechanisms of BBB development in naïve ECs. hPSCs are an attractive model system to complement *in vivo* animal studies because they (i) are human, (ii) permit investigation of developmental processes in contrast to primary or immortalized cells, (iii) are highly scalable, (iv) can be derived from patients to facilitate disease modeling and autologous coculture systems, and (v) are genetically tractable. While widely used hPSC-based BBB models are useful for measuring molecular permeabilities and have been employed to understand genetic contributions to barrier dysfunction (Vatine et al., 2016, 2019; Lim et al., 2017), they have not been shown to proceed through a definitive endothelial progenitor intermediate (Lippmann et al., 2012; Lu et al., 2021) and express epithelial-associated genes (Qian et al., 2017; Delsing et al., 2018; Vatine et al., 2019; Lu et al., 2021). Thus, new models with developmentally relevant differentiation trajectories and definitive endothelial phenotype are needed for improved understanding of developmental mechanisms. Motivated in part by prior use of endothelial cells derived from hematopoietic progenitors in human cord blood to generate BBB models (Boyer-Di Ponio et al., 2014; Cecchelli et al., 2014), we and others recently showed that hPSC-derived naïve endothelial progenitors or ECs are good candidates for such a system (Praça et al., 2019; Nishihara et al., 2020; Roudnicky et al., 2020a, 2020b). For example, Praça *et al*. showed that a combination of VEGF, Wnt3a, and retinoic acid directed EPCs to brain capillary-like ECs with moderate transendothelial electrical resistance (TEER) of ∼60×cm^2^. We previously showed that BBB-like paracellular barrier characteristics are induced in hPSC-EPC-derived ECs after extended culture in a minimal medium. These so-called EECM-BMEC-like cells had TEER and small molecule permeability similar to primary human brain ECs, well-developed tight junctions, and an immune cell adhesion molecule profile similar to brain ECs *in vivo* (Nishihara et al., 2020). In this study, we showed it was possible to use the small molecule Wnt agonist CHIR to induce additional hallmarks of CNS EC phenotype in hPSC-EPC-derived ECs, including canonical GLUT-1, claudin-5, and PLVAP effects (both Passage 1 and 3 CHIR-treated ECs). However, it is important to note that despite the improvements in CNS EC character with CHIR treatment, further improvements to functional endocytosis, and efflux transporter and solute carrier phenotype should be targets of future study and may be facilitated by cocultures and/or additional molecular factors. Along these lines, the Passage 1 CHIR-treated CNS-like ECs would be at a differentiation stage well suited to investigate cues subsequent to Wnt signaling that may be key for the induction of additional CNS EC properties. Alternatively, the Passage 3 CHIR-treated CNS-like ECs may be suitable for other BBB modeling applications. In summary, our work has defined the EC response to Wnt activation in a simplified, human system and established a new hPSC-derived *in vitro* model that will facilitate improved understanding of endothelial barriergenesis.

## MATERIALS AND METHODS

### Key resources table

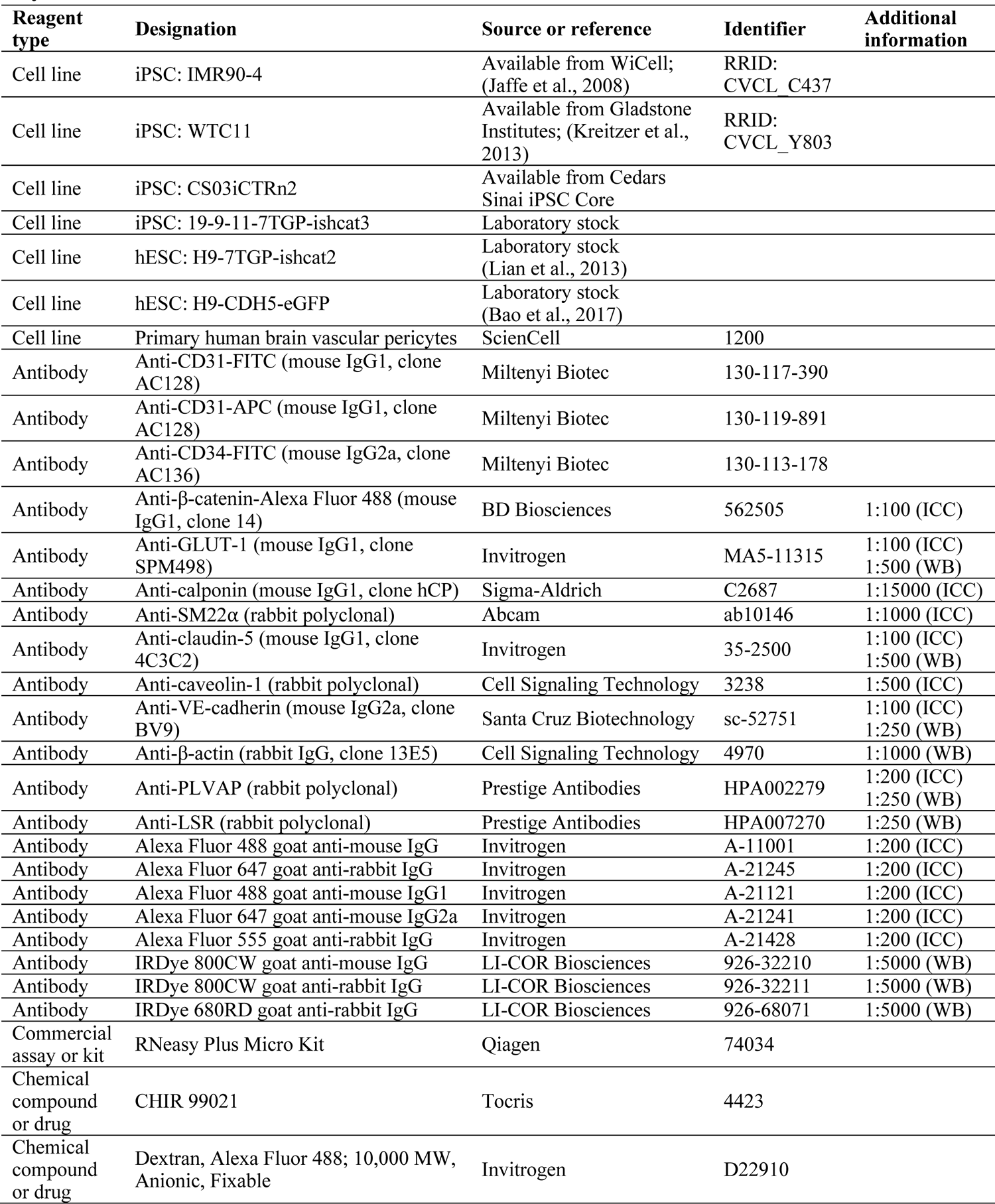
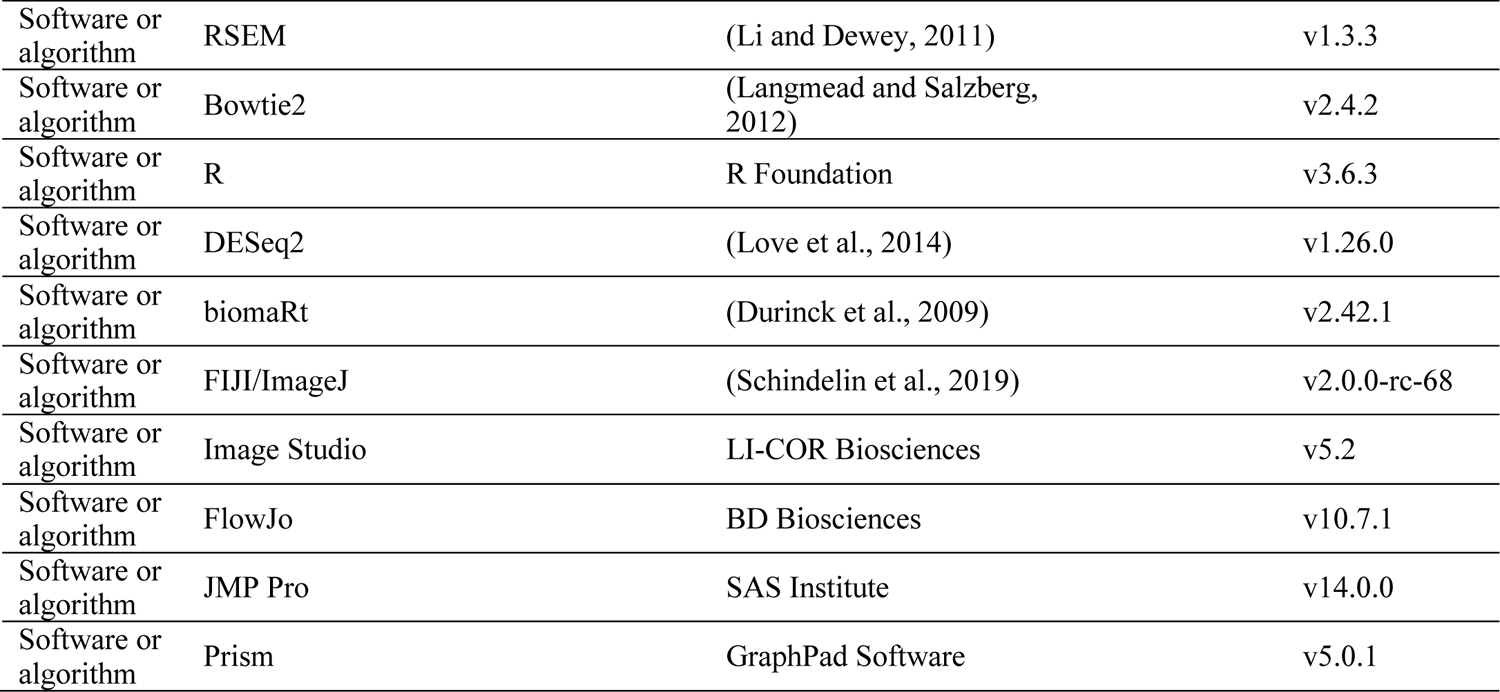

### hPSC maintenance

Tissue culture plates were coated with Matrigel, Growth Factor Reduced (Corning, Glendale, AZ). A 2.5 mg aliquot of Matrigel was thawed and resuspended in 30 mL DMEM/F-12 (Life Technologies, Carlsbad, CA), and the resulting solution used to coat plates at 8.7 µg/cm^2^ (1 mL per well for 6-well plates; 0.5 mL per well for 12-well plates). Plates were incubated at 37°C for at least 1 h prior to use. hPSCs were maintained on Matrigel-coated plates in E8 medium (STEMCELL Technologies, Vancouver, Canada) at 37°C, 5% CO_2_. hPSC lines used were: IMR90-4 iPSC, WTC11 iPSC, CS03iCTRn2 iPSC, H9-CDH5-eGFP hESC, H9-7TGP-ishcat2 hESC, and 19-9-11-7TGP-ishcat3 iPSC. Medium was changed daily. When hPSC colonies began to touch, typically at approximately 70–80% confluence, cells were passaged using Versene (Life Technologies). Briefly, cells were washed once with Versene, then incubated with Versene for 7 min at 37°C. Versene was removed and cells were dissociated into colonies by gentle spraying with E8 medium. Cells were transferred at a split ratio of 1:12 to a new Matrigel-coated plate containing E8 medium.

### Endothelial progenitor cell differentiation

EPCs were differentiated according previously published protocols (Lian et al., 2014; Bao et al., 2016; Nishihara et al., 2020) with slight modifications. On day-3 (D-3), hPSCs were treated with Accutase (Innovative Cell Technologies, San Diego, CA) for 7 min at 37°C. The resulting single cell suspension was transferred to 4× volume of DMEM/F-12 (Life Technologies) and centrifuged for 5 min, 200×g. Cell number was quantified using a hemocytometer. Cells were resuspended in E8 medium supplemented with 10 µM ROCK inhibitor Y-27632 dihydrochloride (Tocris, Bristol, United Kingdom) and seeded on Matrigel-coated 12-well plates at a density of (1.5–2.5)×10^4^ cells/cm^2^, 1 mL per well. Cells were maintained at 37°C, 5% CO_2_. On the following two days (D-2 and D-1), the medium was replaced with E8 medium. The following day (D0), differentiation was initiated by changing the medium to LaSR medium (Advanced DMEM/F-12 [Life Technologies], 2.5 mM GlutaMAX [Life Technologies], and 60 µg/ml L-ascorbic acid 2-phosphate magnesium [Sigma-Aldrich, St. Louis, MO]) supplemented with 7–8 µM CHIR 99021 (Tocris), 2 mL per well. The following day (D1), medium was replaced with LaSR medium supplemented with 7–8 µM CHIR 99021, 2 mL per well. On the following three days (D2, D3, and D4), the medium was replaced with pre-warmed LaSR medium (without CHIR), 2 mL per well.

On D5, EPCs were isolated using CD31 magnetic activated cell sorting (MACS). Cells were treated with Accutase for 15–20 min at 37°C. The resulting cell suspension was passed through a 40 µm cell strainer into an equal volume of DMEM (Life Technologies) supplemented with 10% FBS (Peak Serum, Wellington, CO) and centrifuged for 5 min, 200×g. Cell number was quantified using a hemocytometer. Cells were resuspended in MACS buffer (Dulbecco’s phosphate buffered saline without Ca and Mg [DPBS; Life Technologies] supplemented with 0.5% bovine serum albumin [Sigma-Aldrich] and 2 mM EDTA [Sigma-Aldrich]) at a concentration of 10^7^ cells per 100 µL. The CD31-FITC antibody (Miltenyi Biotec, Auburn, CA) was added to the cell suspension at a dilution of 1:50. The cell suspension was incubated for 30 min at room temperature, protected from light. The cell suspension was brought to a volume of 15 mL with MACS buffer and centrifuged for 5 min, 200×g. The supernatant was aspirated and the pellet resuspended in MACS buffer at a concentration of 10^7^ cells per 100 µL. The FITC Selection Cocktail from the EasySep Human FITC Positive Selection Kit (STEMCELL Technologies) was added at a dilution of 1:10 and the cell suspension was incubated for 20 min at room temperature, protected from light. The Dextran RapidSpheres (magnetic particles) solution from the Selection Kit was added at a dilution of 1:20 and the cell suspension was incubated for an additional 15 min at room temperature.

The cell suspension was brought to a total volume of 2.5 mL with MACS buffer (for total cell number less than 2×10^8^, the approximate maximum yield from two 12-well plates; for a larger number of plates/cells, a total volume of 5 mL was used). 2.5 mL of cell suspension was transferred to a sterile 5 mL round-bottom flow cytometry tube and placed in the EasySep magnet (STEMCELL Technologies) for 5 min. The magnet was inverted to pour off the supernatant, the flow tube removed, the retained cells resuspended in 2.5 mL of MACS buffer, and the flow tube placed back in the magnet for 5 min. This step was repeated 3 times, and the resulting cell suspension transferred to a centrifuge tube, and centrifuged for 5 min, 200×g. Cell number was quantified using a hemocytometer. Resulting EPCs were used directly for experiments as described below or cryopreserved in hECSR medium supplemented with 30% FBS and 10% DMSO for later use. hESCR medium is Human endothelial serum-free medium (Life Technologies) supplemented with 1× B-27 supplement (Life Technologies) and 20 ng/mL FGF2 (Waisman Biomanufacturing, Madison, WI).

### Neural rosette differentiation

Neural rosettes were differentiated according to a previously published protocol (Lippmann et al., 2014) with slight modifications. On D-1, IMR90-4 hPSCs were dissociated with Accutase and seeded on Matrigel-coated plates in E8 medium supplemented with ROCK inhibitor as described above, except the cell seeding density was 5×10^5^ cells/cm^2^. The following day (D0), medium was replaced with E6 medium (DMEM/F-12 supplemented with 64 mg/L L-ascorbic acid 2-phosphate magnesium, 14 µg/L sodium selenium, 543 mg/L sodium bicarbonate, mg/L insulin [Roche, Penzberg, Germany], and 10.7 mg/L holo-transferrin [Sigma-Aldrich]) prepared according to (Chen et al., 2011). Medium was replaced daily with E6 medium on D1 through D5. On D6, medium was replaced with hECSR medium lacking FGF2. The following day (D7), the resulting neural rosette-conditioned medium (NR-CM) was harvested and stored at 4°C, and fresh hECSR medium lacking FGF2 was replaced. NR-CM was likewise harvested on D8, D9, and D10. The resulting NR-CM aliquots were pooled, passed through a 0.2 µm filter, supplemented with 20 ng/mL FGF2, and used for experiments as described below.

### Astrocyte differentiation

Astrocytes were differentiated via an hPSC-derived EZ sphere intermediate according to previously published protocols (Ebert et al., 2013; Sareen et al., 2014; Canfield et al., 2017). Briefly, CS03iCTRn2 hPSCs were dissociated with Versene and colonies were transferred to an ultra-low attachment T-25 flask containing EZ sphere culture medium (a mixture of DMEM and F-12 medium in a 7:3 ratio supplemented with 1× B-27 supplement minus vitamin A [Life Technologies], 2 µg/mL heparin [Sigma-Aldrich], 100 ng/mL EGF [Peprotech], 100 ng/mL FGF2, and 1× antibiotic-antimycotic [Life Technologies]). Half of the volume of EZ sphere culture medium was replaced on Mondays, Wednesdays, and Fridays. EZ spheres were passaged every Friday by mechanical dissociation with 2–4 passes on a McIlwain Tissue Chopper (Campden Instruments, Loughborough, United Kingdom), with half of the resulting aggregates returned to the flask and half discarded. To convert EZ spheres into astrospheres, which are neural stem cell aggregates with enhanced astrocyte differentiation potential, medium was changed to DMEM/F-12 supplemented with 1× N-2 supplement (Life Technologies), 2 µg/mL heparin, 1× MEM-non-essential amino acids solution (Life Technologies), and 0.5 µM all-trans retinoic acid (Sigma-Aldrich) and replaced daily for 11 days. The resulting spheres were passaged as described above and returned to EZ sphere culture medium, which was replaced on Mondays, Wednesdays, and Fridays. Astrospheres were passaged on Fridays as described above and cultured for at least 30 passages prior to initiating astrocyte differentiation. To differentiate astrocytes, astrospheres were treated with Accutase for 10–15 min at 37°C, followed by gentle pipetting to dissociate and singularize the cells. The resulting single cell suspension was transferred to 4× volume of DMEM/F-12 and centrifuged for 5 min, 200×g. Cell number was quantified using a hemocytometer. Cells were resuspended in EZ sphere culture medium and seeded on Matrigel-coated plates at approximately 2.5×10^4^ cells/cm^2^. The following day, medium was changed to astrocyte differentiation medium (DMEM/F-12 supplemented with 1× N-2 supplement, 2 µg/mL heparin, and 1× MEM-non-essential amino acids solution). This medium was replaced every other day for 2 weeks. Medium was then replaced with hECSR medium lacking FGF2. The following day, the resulting astrocyte-conditioned medium (Astro-CM) was harvested and stored at 4°C, and fresh hECSR medium lacking FGF2 was replaced. Astro-CM was likewise harvested on the following 3 days. The resulting Astro-CM aliquots were pooled, passed through a 0.2 µm filter, supplemented with 20 ng/mL FGF2, and used for experiments as described below.

### Endothelial cell culture and treatment

Collagen IV (Sigma-Aldrich) was dissolved in 0.5 mg/mL acetic acid to a final concentration of 1 mg/mL. Collagen IV-coated plates were prepared by diluting a volume of this stock solution 1:100 in water, adding the resulting solution to tissue culture plates, or #1.5 glass bottom plates (Cellvis, Sunnyvale, CA) for cells intended for confocal imaging (1 mL per well for 6-well plates, 0.5 mL per well for 12-well plates, 0.25 mL per well for 24-well plates), and incubating the plates for 1 h at RT. Collagen IV coating solution was removed and EPCs obtained as described above were suspended in hECSR medium and plated at approximately 3×10^4^ cells/cm^2^. In some experiments, cells were suspended in NR-CM, Astro-CM, or Peri-CM. In some experiments, ligands and small molecules were added to hECSR medium or Peri-CM: CHIR 99021 (Tocris) was used at 4 µM except where indicated; DMSO (Sigma-Aldrich) was used as a vehicle control for CHIR; Wnt3a (R&D Systems) was used at 20 ng/mL; Wnt7a (Peprotech, Rocky Hill, NJ) was used at 50 ng/mL; Wnt7b (Abnova, Tapei, Taiwan) was used at 50 ng/mL; R-spondin 1 (Rspo1; Peprotech) was used at 50 ng/mL; doxycycline was used at 1, 2, or 4 µg/mL. The hECSR medium or CM, including any ligands or small molecules, was replaced every other day until confluent (typically 6 days). We denote this time point “Passage 1.”

For extended culture, ECs were selectively dissociated and replated as previously described (Nishihara et al., 2020). Cells were incubated with Accutase until endothelial cells appeared round, typically 2–3 min at 37°C. The plate was tapped to release the ECs while SMLCs remained attached, and the EC-enriched cell suspension transferred to 4× volume of DMEM/F-12 and centrifuged for 5 min, 200×g. Cells were resuspended in hECSR medium and seeded on a new collagen IV-coated plate at approximately 3×10^4^ cells/cm^2^. hECSR medium was replaced every other day until confluent (typically 6 days). The selective dissociation and seeding described above was repeated, and hECSR medium was again replaced every other day until confluent (typically 6 days). We denote this time point “Passage 3.” In one experiment, these steps were repeated for another two passages. Except where indicated, CHIR 99021 or vehicle (DMSO) was included in the hECSR medium for the entire duration of culture.

### RNA-seq

RNA-seq was performed on ECs and SMLCs from the IMR90-4 hPSC line. Four independent differentiations were performed, with DMSO-, CHIR-, and Wnt7a/b-treated ECs at Passage 1 analyzed from all four differentiations. DMSO- and CHIR-treated ECs at Passage 3 were analyzed from three of the four differentiations. DMSO-treated SMLCs at Passage 1 were analyzed from two of the four differentiations. Fluorescence-activated cell sorting (FACS) was used to isolate CD31^+^ ECs and CD31^-^ SMLCs from mixed Passage 1 cultures. Cells were incubated with Accutase for 10 min at 37°C, passed through 40 µm cell strainers into 4× volume of DMEM/F-12, and centrifuged for 5 min, 200×g. Cells were resuspended in MACS buffer and incubated with CD31-APC antibody (Miltenyi Biotec) for 30 min at 4°C, protected from light. The cell suspension was brought to a volume of 15 mL with MACS buffer and centrifuged at 4°C for 5 min, 200×g. Cells were resuspended in MACS buffer containing 2 µg/mL 4’,6-diamidino-2-phenylindole (DAPI; Life Technologies). A BD FACSAria III Cell Sorter (BD Biosciences, San Jose, CA) was used to isolate DAPI^-^CD31^+^ cells (live ECs) and DAPI^-^CD31^-^ cells (live SMLCs). The resulting cell suspensions were centrifuged at 4°C for 5 min, 200×g, and cell pellets immediately processed for RNA extraction as described below.

RNA was isolated using the RNeasy Plus Micro Kit (Qiagen, Germantown, MD). Buffer RLT Plus supplemented with 1% β-mercaptoethanol was used to lyse cells (pellets from FACS of Passage 1 cells, or directly on plates for Passage 3 ECs). Lysates were passed through gDNA Eliminator spin columns, loaded onto RNeasy MinElute spin columns, washed with provided buffers according to manufacturer instructions, and eluted with RNase-free water. Sample concentrations were determined using a NanoDrop spectrophotometer (Thermo Scientific, Waltham, MA) and RNA quality assayed using an Agilent 2100 Bioanalyzer with Agilent RNA 6000 Pico Kit (Agilent, Santa Clara, CA). First-strand cDNA synthesis was performed using the SMART-Seq v4 Ultra Low Input RNA kit (Takara Bio, Mountain View, CA) with 5 ng input RNA followed by 9 cycles of PCR amplification and library preparation using the Nextera XT DNA Library Prep Kit (Illumina, San Diego, CA). Sequencing was performed on a NovaSeq 6000 (Illumina), with approximately 40–60 million 150 bp paired-end reads obtained for each sample.

FASTQ files were aligned to the human genome (hg38) and transcript abundances quantified using RSEM (v1.3.3) (Li and Dewey, 2011) calling bowtie2 (v2.4.2) (Langmead and Salzberg, 2012). Estimated counts from RSEM were input to DESeq2 (v1.26.0) (Love et al., 2014) implemented in R (v3.6.3) for differential expression analysis. Elsewhere, transcript abundances are presented as transcripts per million (TPM). Differentiation pairing as described above was included in the DESeq2 designs. The Wald test with Benjamini-Hochberg correction was used to generate adjusted P-values. Principal component analysis was performed on counts after the DESeq2 variance stabilizing transformation. Bulk RNA-seq data from the literature (FASTQ files; see *Previously published datasets used*) were obtained from the Gene Expression Omnibus (GEO). These FASTQ files were aligned to the mouse genome (mm10) and transcript abundances quantified as described above. DESeq2 was used for differential expression analysis as described above. For direct comparison of human and mouse data, the biomaRt package (v2.42.1) (Durinck et al., 2009) and Ensembl database (Yates et al., 2019) was used to map human gene names to mouse homologs. Venn diagrams were generated using the tool available at http://bioinformatics.psb.ugent.be/webtools/Venn/. To identify solute carrier and efflux transporter genes highly expressed at the human BBB *in vivo*, we used five human brain scRNA-seq datasets (see *Previously published datasets used*) integrated in a previous meta-analysis (Gastfriend et al., 2021). *SLC* and *ABC* genes with average expression greater than 100 TPM in endothelial cells across the five independent datasets were selected.

### Immunocytochemistry

Immunocytochemistry was performed in 24-well plates. Cells were washed once with 500 µL DPBS and fixed with 500 µL cold (–20°C) methanol for 5 min, except cells intended for calponin/SM22a detection, which were fixed with 500 µL of 4% paraformaldehyde for 15 min. Cells were washed three times with 500 µL DPBS and blocked in 150 µL DPBS supplemented with 10% goat serum (Life Technologies) for 1 h at room temperature, except cells intended for calponin/SM22⍺ detection, which were blocked and permeabilized in DPBS supplemented with 3% BSA and 0.1% Triton X-100. Primary antibodies diluted in 150 µL of the above blocking solutions (see *Key Resources Table* for antibody information) were added to cells and incubated overnight at 4°C on a rocking platform. Cells were washed three times with 500 µL DPBS. Secondary antibodies diluted in 150 µL of the above blocking solutions (see *Key Resources Table* for antibody information) were added to cells and incubated for 1 h at room temperature on a rocking platform, protected from light. Cells were washed three times with 500 µL DPBS, followed by 5 min incubation with 500 µL DPBS plus 4 µM Hoechst 33342 (Life Technologies). Images were acquired using an Eclipse Ti2-E epifluorescence microscope (Nikon, Tokyo, Japan) with a 20× objective or an A1R-Si+ confocal microscope (Nikon) with a 100× oil objective. Confocal images were acquired with 1 µm slice spacing.

Images were analyzed using FIJI (ImageJ) software. For epifluorescence images, 5 fields (20×) were analyzed per well, with 3–4 wells per treatment condition. For quantification of cell number, EC colonies were manually outlined, and the Analyze Particles function was used to estimate the number of nuclei within the EC colonies. Nuclei outside the EC colonies were manually counted. EC purity (% EC) was calculated as the number of nuclei within EC colonies relative to total nuclei. To estimate % GLUT-1^+^ ECs, cells within the EC colonies with membrane-localized GLUT-1 immunoreactivity (e.g., arrowheads in Figure 2B) were manually counted. For quantification of fluorescence intensity in epifluorescence images, EC colonies were manually outlined, and the Measure function was used to obtain the mean fluorescence intensity for each image channel (fluorophore). A cell-free area of the plate was similarly quantified for background subtraction. Following background subtraction, the mean fluorescence intensity of each protein of interest was normalized to the mean fluorescence intensity of Hoechst to correct for effects of cell density. For confocal images, 3–4 fields (100×) containing only VE-cadherin^+^ ECs were analyzed per well, with 4 wells per treatment condition. The first slice with visible nuclei (closest to glass) was defined as *Z* = 0, and the Measure function was used to obtain the mean fluorescence intensity for each image channel (fluorophore) in each slice from *Z* = 0 to *Z* = 7 µm. A cell-free area of the plate was similarly quantified for background subtraction. After background subtraction, to approximate total abundance (area under the fluorescence versus *Z* curve, AUC) for each channel, mean fluorescence intensities were summed across all slices. AUC for the proteins of interest were normalized to Hoechst AUC.

### Western blotting

To enrich samples from Passage 1 cultures for ECs, the Accutase-based selective dissociation method described above was employed. Dissociated cells were centrifuged for 5 min, 200×g, and resulting cell pellets were lysed in RIPA buffer (Rockland Immunochemicals, Pottstown, PA) supplemented with 1× Halt Protease Inhibitor Cocktail (Thermo Scientific). Passage 3 cells were lysed with the above buffer directly on plates. Lysates were centrifuged at 4°C for 5 min, 14,000×g, and protein concentration in supernatants quantified using the Pierce BCA Protein Assay Kit (Thermo Scientific). Equal amounts of protein were diluted to equal volume with water, mixed with sample buffer, and heated at 95°C for 5 min, except lysates intended for GLUT-1 Western blotting, which were not heated. Samples were resolved on 4– 12% Tris-Glycine gels and transferred to nitrocellulose membranes. Membranes were blocked for 1 h in tris-buffered saline plus 0.1% Tween-20 (TBST) supplemented with 5% non-fat dry milk. Primary antibodies (see *Key Resources Table* for antibody information) diluted in TBST plus 5% non-fat dry milk were added to membranes and incubated overnight at 4°C on a rocking platform. Membranes were washed five times with TBST. Secondary antibodies (see *Key Resources Table* for antibody information) diluted in TBST were added to membranes and incubated for 1 h at room temperature on a rocking platform, protected from light. Membranes were washed five times with TBST and imaged using an Odyssey 9120 (LI-COR, Lincoln, NE). Band intensities were quantified using Image Studio software (LI-COR).

### Dextran accumulation assay

A fixable, Alexa Fluor 488-conjugated dextran with an average molecular weight of 10 kDa (Invitrogen) was used as a tracer to estimate total fluid-phase endocytosis. Dextran was added at 10 µM to the medium of Passage 1 cultures. Plates were incubated on rotating platforms at 37°C or 4°C for 2 h. Medium was removed and cells were washed once with DPBS, and then incubated with Accutase for 10 min at 37°C. Cell suspensions were passed through 40 µm cell strainers into 4× volume of DMEM/F-12 and centrifuged for 5 min, 200×g. Cells were resuspended in MACS buffer and incubated with the CD31-APC antibody (Miltenyi Biotec) for 30 min at 4°C, protected from light. Cell suspensions were brought to a volume of 5 mL with MACS buffer and centrifuged at 4°C for 5 min, 200×g. Pellets were resuspended in DPBS supplemented with 4% paraformaldehyde and incubated for 15 min at room temperature, protected from light. Cells were centrifuged for 5 min, 200×g. Pellets were resuspended in MACS buffer and analyzed on a BD FACSCalibur flow cytometer (BD Biosciences). FlowJo software (BD Biosciences) was used to gate CD31^+^ cells and quantify geometric mean fluorescence intensity and coefficient of variation (CV) of dextran.

### Statistics

Individual wells of cultured cells that underwent identical experimental treatments are defined as replicates, and all key experiments were repeated using multiple independent hPSC differentiations. Detailed information about replication strategy is provided in figure legends. Student’s *t* test was used for comparison of means from two experimental groups. One-way analysis of variance (ANOVA) was used for comparison of means from three or more experimental groups, followed by Dunnett’s post-hoc test for comparison of multiple treatments to a single control, or Tukey’s honest significant difference (HSD) post-hoc test for multiple pairwise comparisons. When data from multiple differentiations were combined, two-way ANOVA (one factor being the experimental treatment and one factor being the differentiation) was used for comparison of means to achieve blocking of differentiation-based variability, followed by post-hoc tests as described above if more than two experimental treatments were compared. For fluorescence intensities (a.u.), two-way ANOVA was performed prior to normalization of these values to the control group within each differentiation (for visualization in plots). Statistical tests were performed in JMP Pro (v15.0.0). For RNA-seq differential expression analysis, the DESeq2 Wald test with Benjamini-Hochberg correction was used to calculate P-values. Descriptions of the statistical tests used are provided in figure legends.

## Acknowledgements

We acknowledge the University of Wisconsin–Madison Biotechnology Center Gene Expression Center and DNA Sequencing Facility for providing library preparation and next generation sequencing services. We acknowledge the University of Wisconsin–Madison Biochemistry Optical Core for use of a confocal microscope. We acknowledge the University of Wisconsin Carbone Cancer Center Flow Cytometry Laboratory (supported by NIH Cancer Center Support Grant P30 CA014520) for performing FACS. This work was funded by NIH (NS103844 and NS107461 to EVS and SPP), the Swiss National Science Foundation (grant 310030_189080 to BE), and the Bern Center for Precision Medicine to BE. BDG was supported by NIH Biotechnology Training Program grant T32 GM008349 and the National Science Foundation Graduate Research Fellowship Program under grant number 1747503. HN was supported by a JSPS Overseas Research Fellowship.

## Supplementary files

• Supplementary file 1. RNA-sequencing gene expression data for hPSC-derived ECs and SMLCs. Abundances are provided in transcripts per million (TPM).

• Supplementary file 2. RNA-sequencing differential expression analysis of hPSC-derived ECs. DESeq2-derived average expression (baseMean), log_2_(fold change), Wald statistic, P-value (Wald test), and adjusted P-value (Benjamini-Hochberg correction) are shown. (A) Passage 1 CHIR-treated ECs versus Passage 1 DMSO-treated ECs. (B) Passage 1 Wnt7a/b-treated ECs versus Passage 1 DMSO-treated ECs. (C) Passage 3 CHIR-treated ECs versus Passage 3 DMSO-treated ECs. (D) Passage 3 DMSO-treated ECs versus Passage 1 DMSO-treated ECs. (E) Lists of upregulated and downregulated genes comprising the intersection of the comparisons in (A) and (B), and (A) and (C), used to generate Venn diagrams in Figure 8E.

• Supplementary file 3. Wnt-regulated EC genes in multiple contexts. (A) Differential expression analysis of P7 murine brain, liver, lung, and kidney ECs (Sabbagh et al., 2018). DESeq2-derived average expression (baseMean), log_2_(fold change), Wald statistic, P-value (Wald test), and adjusted P-value (Benjamini-Hochberg correction) are shown. (B) Lists of brain-enriched and brain-depleted genes comprising the intersection of the comparisons in (A). (C-E) Differential expression analysis of adult murine ECs with β-catenin-stabilization versus controls from pituitary (Wang et al., 2019) (C), liver (Munji et al., 2019) (D), and brain ECs cultured *in vitro* (Sabbagh and Nathans, 2020) (E). DESeq2-derived average expression (baseMean), log_2_(fold change), Wald statistic, P-value (Wald test), and adjusted P-value (Benjamini-Hochberg correction) are shown. (F) Lists of concordantly Wnt-regulated genes in Passage 1 hPSC-derived ECs and the three comparisons shown in (C-E), from the set of brain-enriched and brain-depleted genes identified in (B).

## Data availability

RNA-seq data have been deposited in GEO.

## Previously published datasets used

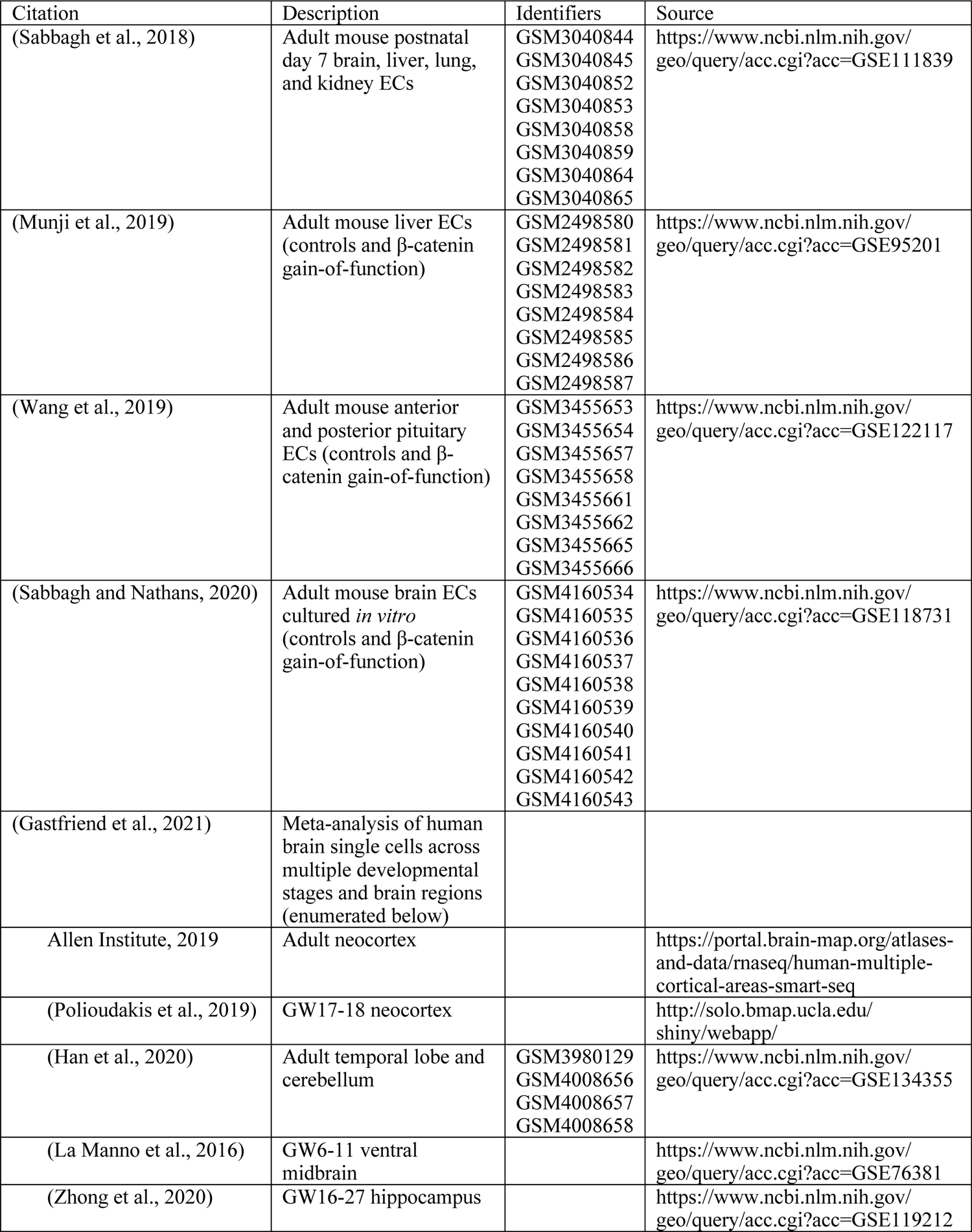

